# Minkowski Tensors as a Lightweight and Interpretable Representation for Three-Dimensional Morphology

**DOI:** 10.64898/2026.09.27.754865

**Authors:** Yajushi Khurana, Keisuke Ishihara

## Abstract

Quantitative analysis of biological shape is central to understanding how form encodes function. Despite rapid advances in volumetric imaging and segmentation, analysis and of three-dimensional morphology remains challenging due to complex geometries and topologies found in biology. Existing approaches either rely on handcrafted metrics that lack generalizability or employ data-driven neural models that sacrifice interpretability and robustness. Here, we introduce Minkowski Tensors (MT), a family of integral-geometric shape descriptors that extend classical scalar measures of volume, surface area, and curvature into tensor-valued quantities encoding direction and anisotropy. MT form a provably complete yet compact and interpretable feature set that captures volumetric and curvature-based information in a transformation-and scale-covariant, topology-aware manner. We systematically derive rotation-invariant and covariant subgroups of the tensor family, explicitly constructing measures for invariance and covariance, together with Euler-characteristic-based subgroups that capture topological features such as holes and connected components. We benchmark MT on diverse segmented 3D biomedical images spanning vascular and adrenal structures from magnetic resonance angiography (MRA) and computed tomography (CT), and dividing cell nuclei imaged by confocal microscopy, demonstrating competitive or superior classification performance relative to classical geometric descriptors and neural baselines while retaining direct biological interpretability. Our results establish Minkowski tensors as a concise yet powerful framework for discriminative analysis of complex 3D biological morphology.

## Introduction

Morphology is a fundamental descriptor of biological state. Across scales from subcellular organelles to tissues and organs, shape reflects underlying biochemical, mechanical, and genetic processes, linking geometry to physiology and pathology (1–4). Advances in three-dimensional (3D) volumetric imaging and segmentation have enabled systematic acquisition of morphological data at unprecedented resolution and throughput, yet obtaining biologically meaningful and generalizable features from these datasets remains a major bottleneck. Compared to two-dimensional (2D) images, 3D shapes exhibit higher degrees of freedom in geometric variation, making it difficult to define robust and interpretable metrics for comparing complex morphologies.

Quantitative morphology hinges on mapping high-dimensional geometry to a compact, comparable representation. One of the most common uses of such representations is to discriminate phenotypes or functional classes, such as healthy versus diseased states or distinct functional biological stages, using geometric and statistical features. Within discriminative analysis, the central question is how 3D shape is represented, and two contrasting strategies have emerged: learned representations, in which deep neural networks infer features directly from data, and explicitly defined representations, in which geometric and topological descriptors are computed from the shape and paired with standard classifiers. Learned representations achieve strong discriminative performance by learning hierarchical features through architectural and data-driven priors, but typically require large amounts of training data and substantial computational resources, and frequently exhibit limited robustness and biological interpretability in the absence of explicit geometric constraints (5). Explicitly defined representations, in contrast, rely on low-dimensional descriptors derived from the underlying geometry and topology of objects to quantify spatial relationships, boundaries, and symmetries, and are typically coupled with relatively simple classifiers (e.g., linear models or support vector machines) so that discriminative performance is driven primarily by the quality of the representation rather than model complexity. By encoding prior knowledge directly at the feature level, explicitly defined representations offer greater transparency, reproducibility, and often improved robustness under limited data or varying acquisition conditions, but critically depend on the choice of features and their ability to capture biologically relevant variation.

The effectiveness of explicitly defined representations therefore hinges on the specific descriptor chosen, yet the most widely used ones each carry significant limitations. Spherical harmonics expansion (SHE) has been widely applied for shape reconstruction due to its invertible representation (6) and has also been integrated into discriminative pipelines (7), but despite its expressive power it is inherently restricted to near-spherical topologies and fails to adequately capture non-zero genus structures (2, 6). CellProfiler (CP), a widely-used open-source image analysis platform with a graphical user interface (GUI) (8), provides interpretable, SciPy- and scikit-image–derived morphological metrics such as size, volume, and surface characteristics for 3D datasets, but these hand-crafted features are limited in their ability to handle multi-body configurations and can become computationally inefficient for large-scale volumetric analysis. Other classical representations, including landmark-based and distance-based descriptors, either require extensive expert input (9) or lack sufficient discriminative power for closely related morphologies (10), respectively. Collectively, these limitations highlight the need for a compact, interpretable feature set that explicitly encodes geometric and topological priors, retaining the robustness and efficiency of classical machine learning while capturing sufficient morphological information to discriminate complex 3D biological structures.

To address this need, we present Minkowski tensors (MT), a morphology descriptor: integral-geometric quantities that extend scalar measures of size and topology to encode directional and geometric information in a transformation-covariant manner. Crucially, Minkowski tensors are not an ad hoc choice — they form a complete family of motion-covariant valuations (Hadwiger–Alesker). An ideal descriptor family of this kind should span both invariant descriptors, for when shape alone matters, and covariant descriptors, for when orientation or size carries biological meaning, while also accommodating the non-zero genus and multi-body structures common in biological datasets. Minkowski tensors satisfy all of these criteria. MT-based approaches have already been applied across several domains, including 2D galaxy arrangements (11), neuronal cell shape (12), and epithelial monolayer orientation (13), and in 3D to the geometric characterization of trabecular bone (14), but the full rank 0-2 tensor family and its derived invariants remain largely unevaluated as a feature set for 3D biological classification. Building on this foundation, we systematically evaluate the full range of Minkowski tensor subgroups—including raw tensors, rotation-invariant reductions, and topology-augmented variants—and perform ablations to identify which components carry the discriminative signal, augmenting with additional tensor-derived, physically interpretable quantities to capture comprehensive anisotropic shape and topological information. We evaluate our feature set with respect to (i) its fidelity as a 3D shape descriptor, including invariance to transformation; (ii) biological interpretability and discriminative performance across diverse 3D biological datasets including multibody and non-zero genus morphologies; and (iii) benchmarking against existing discriminative approaches, thereby advancing the quantitative analysis of morphology in complex biological systems.

## Materials and Methods

### Imaging Datasets

#### MedMNIST 3D Biomedical Imaging Dataset

We evaluated Minkowski descriptors on the Adrenal3D and Vessel3D subsets of MedMNIST v2 (15), volumetric segmentations derived from CT and MRA scans respectively, each comprising approximately 10^3^ samples at two spatial resolutions (28^3^ and 64^3^) with a standardized 7:2:1 train/test/validation split. Adrenal3D contains binary classifications of normal versus hyperplastic adrenal glands; Vessel3D contains healthy versus aneurysmal vessels; in addition, Adrenal3D samples carry left/right annotations from the source repository (15), which we use to assess orientation sensitivity.

#### Allen Cell Institute hiPSC Dataset

To assess performance on topologically complex morphologies, we evaluated MT on manually annotated 3D nuclear segmentation masks from the Allen Cell Institute hiPSC dataset (2), comprising 5606 nuclei distributed across six cell-cycle stages (Interphase, Prophase, Early Prometaphase, Prometaphase/Metaphase, and two Anaphase/Telophase/Cytokinesis stages) with a strongly imbalanced class distribution. Nuclear morphologies exhibit substantial variability in shape, include multi-body configurations, and span a wide range of topological complexity including non-zero genus structures; we applied the same stratified 7:2:1 split as for MedMNIST (see Supplementary Methods for details).

### Calculation of Shape Descriptors

#### Translational Alignment and Preprocessing

Prior to descriptor computation, all segmentation masks were translated so that the center of mass coincided with the array center, placing each object in a common geometric reference frame shared by all descriptor methods. The hiPSC nuclei additionally required downsampling and center-padding to a fixed 96 × 256 × 256 array to provide fixed-size inputs for deep-learning baselines while preserving intrinsic shape information; see Supplementary Methods for full details.

#### Minkowski Tensors

We computed Minkowski tensors at ranks 0–2 using pykarambola (v0.5.1) (16), yielding four scalar functionals, four rank-1 vectors, and six rank-2 tensors together with derived rotation-invariant quantities (eigenvalues, anisotropy indices *β*, traces, determinants, and trace ratios); the physical interpretation and transformation properties of each quantity are summarized in Table 1, and mathematical definitions are given in Supplementary Methods. We systematically evaluated nested and overlapping feature subgroups spanning the raw tensorial components, rotation-invariant reductions, augmented combinations, and a 36-dimensional full invariant set (see Supplementary Methods for the full ablation design).

**Table 1.** Minkowski tensors and derived quantities used in this work, with their physical interpretation and their behavior under rotation and scaling, sensitivity to topology, and additivity. The rotation and scale columns are three-valued: *inv*. (invariant, the value is unchanged), *cov*. (covariant, the value transforms predictably with the object—for scaling, by a fixed power of the scale factor). Translation is omitted, as all objects are translationally aligned prior to computation (Translational Alignment, Section). *Topology* marks quantities derived from the *W*_3_ (Gaussian-curvature) integral that encode the Euler characteristic, and are thus sensitive to topological features such as holes and connected components. *Additive* marks quantities satisfying the inclusion–exclusion (valuation) property; ratios, spectral quantities, and anisotropy indices are not additive.

| Quantity | Rank | Interpretation | Rotation | Scale | Topology | Additive |
| --- | --- | --- | --- | --- | --- | --- |
| *Raw valuations* |  |  |  |  |  |  |
| $W_0$ (w000) | 0 | Volume | inv. | cov. | × | ✓ |
| $W_1$ (w100) | 0 | Surface area | inv. | cov. | × | ✓ |
| $W_2$ (w200) | 0 | Integral mean curvature | inv. | cov. | × | ✓ |
| $W_3$ (w300) | 0 | Euler characteristic (Gauss–Bonnet) | inv. | cov. | ✓ | ✓ |
| $W_0^{1,0}$ (w010) | 1 | Volume moment; $W_0^{1,0}/W_0$ = center of mass | cov. | cov. | × | ✓ |
| $W_1^{1,0}$ (w110) | 1 | Surface moment; $W_1^{1,0}/W_1$ = area-weighted centroid | cov. | cov. | × | ✓ |
| $W_2^{1,0}$ (w210) | 1 | Curvature moment; $W_2^{1,0}/W_2$ = curvature-weighted centroid | cov. | cov. | × | ✓ |
| $W_3^{1,0}$ (w310) | 1 | Gaussian-curvature moment; $W_3^{1,0}/W_3$ = topology-weighted centroid | cov. | cov. | ✓ | ✓ |
| $W_0^{2,0}$ (w020) | 2 | Solid moment tensor | cov. | cov. | × | ✓ |
| $W_1^{2,0}$ (w120) | 2 | Hollow moment tensor | cov. | cov. | × | ✓ |
| $W_2^{2,0}$ (w220) | 2 | Wire moment tensor | cov. | cov. | × | ✓ |
| $W_3^{2,0}$ (w320) | 2 | Vertex moment tensor | cov. | cov. | ✓ | ✓ |
| $W_1^{0,2}$ (w102) | 2 | Normal distribution | cov. | cov. | × | ✓ |
| $W_2^{0,2}$ (w202) | 2 | Curvature distribution | cov. | cov. | × | ✓ |
| *Derived quantities (per rank-2 tensor)* |  |  |  |  |  |  |
| Eigenvalues $\xi_{1,2,3}$ | — | Principal-axis magnitudes of the tensor | inv. | cov. | × | × |
| Trace | — | Sum of eigenvalues; isotropic tensor concentration | inv. | cov. | × | ✓ |
| Determinant | — | Product of eigenvalues | inv. | cov. | × | × |
| Anisotropy index $\beta$ | — | $\xi_{\min}/\xi_{\max}$; tensor isotropy | inv. | inv. | × | × |
| Trace ratio | — | Ratio of traces; tensor concentration per scalar | inv. | cov. | × | × |

#### CellProfiler Features

CellProfiler (8) features were extracted using the `MeasureSizeShape` module and evaluated as several subgroups: the full feature set, shape features only, location features only, and individual interpretable descriptors (surface area, solidity, and their combination).

#### Spherical Harmonic Descriptors

Spherical harmonic descriptors were computed using `aics-shparam` (2); we evaluated expansions from *l*_max_ =1 to *l*_max_ = 16, as performance plateaued at low orders and declined at higher orders consistent with high-order coefficients encoding increasingly noise-dominated detail.

### Experiments

***Classification Task and Benchmarking***. Each feature set was standardized and evaluated with an SVM classifier (17) tuned by Bayesian optimization, and repeated with a LightGBM classifier (18) to confirm that findings reflect the feature set rather than classifier choice. Performance was measured by balanced accuracy (BA) and geometric mean (G-mean), both appropriate for imbalanced data; the complete procedure was repeated three times with independent random seeds and results are reported as mean ± standard deviation. We benchmarked against CellProfiler features, spherical harmonic descriptors, and deep-learning baselines (published ResNet-18/50 results for MedMNIST (15); trained 3D ResNet-34/50/101 for hiPSC nuclei (2)). See Supplementary Methods for full pipeline and benchmarking details.

#### Importance Analysis

To identify which individual descriptors drive classification, we performed permutation feature importance analysis (19), quantifying the decrease in balanced accuracy when each feature’s values are randomly permuted while all others are held fixed. We reported grouped permutation importance, permuting strongly correlated features (|*r*| *>* 0.9) jointly to avoid underestimating collinear descriptors; see Supplementary Methods for details.

#### Trajectory Analysis and Statistical Tests for Biological Interpretability

To probe biological interpretability, we characterized the distributions of top-ranked features across classes using rank-based statistical tests (Mann–Whitney, Kruskal–Wallis with Dunn post-hoc correction, effect sizes via Cliff’s *δ* and Cohen’s *d*; see Supplementary Methods for full statistical details). We further visualized the global structure of the Minkowski feature space using PHATE (20) to test whether biologically adjacent cell-cycle stages occupy adjacent regions in feature space.

#### Rotational Robustness

During benchmarking on Vessel3D, CellProfiler outperformed Minkowski tensor features, and permutation importance analysis revealed that bounding-box extent features along the primary imaging axis (*X*-direction: BoundingBoxMinimum/Maximum X) ranked among the top CP discriminators in standard orientation—indicating a systematic directional bias in the dataset rather than intrinsic shape signal. To test whether this advantage reflected genuine shape information or merely consistent orientations, we created a randomized-orientation variant of Vessel3D by applying an independent rotation drawn uniformly from *SO*(3) to each object mask, recomputing all features on the rotated masks, and evaluating the identical classification pipeline. This rotational robustness experiment is applied to Vessel3D only, as it was the dataset where such orientation-dependent CP features dominated.

### Availability and Implementation

The MedMNIST v2 dataset is available at https://medmnist.com/ (Zenodo: https://doi.org/10.5281/zenodo.10519652) and can be imported using their python package `medmnist`. The human iPSC nuclear segmentation masks are from the Allen Cell WTC-11 hiPSC Single-Cell Image Dataset (2), available from the Allen Institute for Cell Science at https://open.quiltdata.com/b/allencell/packages/aics/mitotic_annotation. Minkowski features were extracted using the pykarambola package (16), available at https://github.com/Ishihara-SynthMorph/pykarambola (pip install pykarambola). The code for this work is available from the lead contact upon request.

## Results

### Covariant descriptors recover organ handedness in adrenal glands

We begin by testing the fidelity of Minkowski tensor descriptors to capture biologically relevant morphological variation and their response to geometric transformations. Many biological problems require capturing an object’s orientation, while others are better served by discarding it entirely. Because the Minkowski tensor family includes both covariant and invariant descriptors, it can accommodate either need. The Adrenal3D dataset illustrates this (see Figure 1A for example meshes of each class). On the Adrenal3D dataset, beyond the diagnostic label (normal versus hyperplastic gland), each sample carries a left/right (L/R) annotation: the adrenal glands occur as bilateral pairs whose two sides are approximate mirror images. This provides a natural test of orientation sensitivity, since L and R glands differ principally in handedness rather than in intrinsic shape. As summarized in Table 1, the raw tensors and vectors are *covariant* under rotation their values transform with the orientation of the object, whereas the scalar functionals and the derived quantities (eigenvalues, anisotropy indices, traces) are rotation-*invariant*. Consequently, if orientation carries information, it should be recoverable from the covariant features alone. These descriptors are computed from binary segmentation masks using pykarambola (Figure 1B), which converts each mask to a triangulated surface mesh and evaluates tensor-valued surface integrals over the mesh faces. Projecting the tensor and vector components onto their first two principal components separates the samples into distinct left and right clusters (Fig. 2A), confirming that these features encode handedness. When the same projection is instead computed from the rotation-invariant measures, the L/R clusters collapse (Fig. 2B), as expected for features that discard orientation. Together these results demonstrate that the descriptor set cleanly separates orientation-dependent from orientation-independent information: covariance is not a limitation but a usable channel, recovering a biologically meaningful axis (bilateral handedness) that invariant descriptors necessarily discard.

**Fig. 1.**
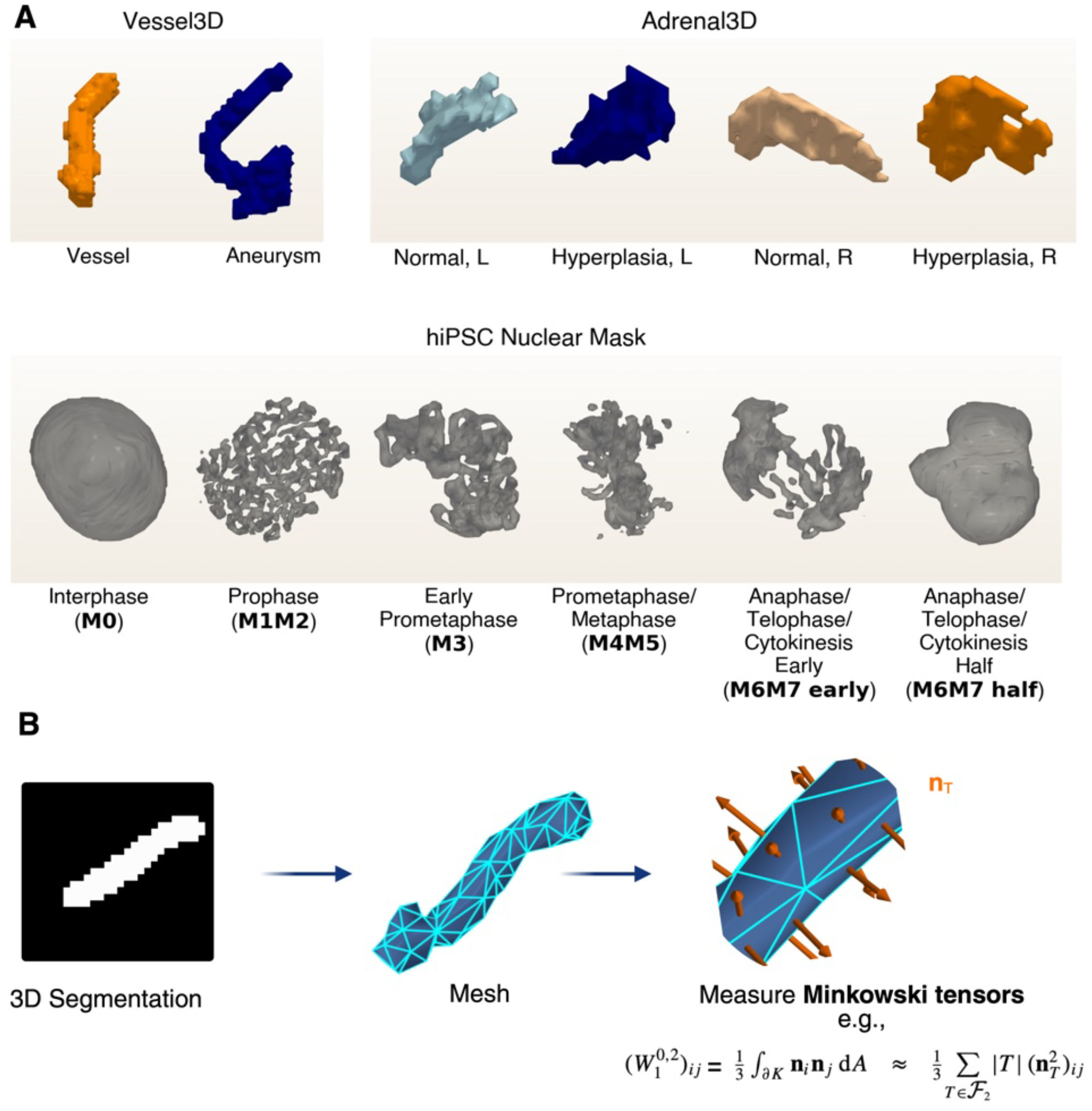
Datasets and Minkowski tensor computation pipeline. (**A**) Representative 3D morphologies from each dataset. *Top:* Vessel3D (healthy vessel, blue; aneurysm, orange) and Adrenal3D segmentation masks from MedMNIST; Adrenal3D has four classes: normal left (light blue), hyperplastic left (dark blue), normal right (light orange), hyperplastic right (dark orange). *Bottom:* hiPSC nuclear masks spanning six cell-cycle stages: Interphase (M0), Prophase (M1M2), Early Prometaphase (M3), Prometaphase/Metaphase (M4M5), Anaphase/Telophase/Cytokinesis Early (M6M7e), and Anaphase/Telophase/Cytokinesis Half (M6M7h). (**B**) Minkowski tensor computation pipeline using pykaram-bola. A binary 3D segmentation is converted to a triangulated surface mesh; tensor-valued surface integrals are then evaluated over the mesh faces to yield the Minkowski feature vector.

**Fig. 2.**
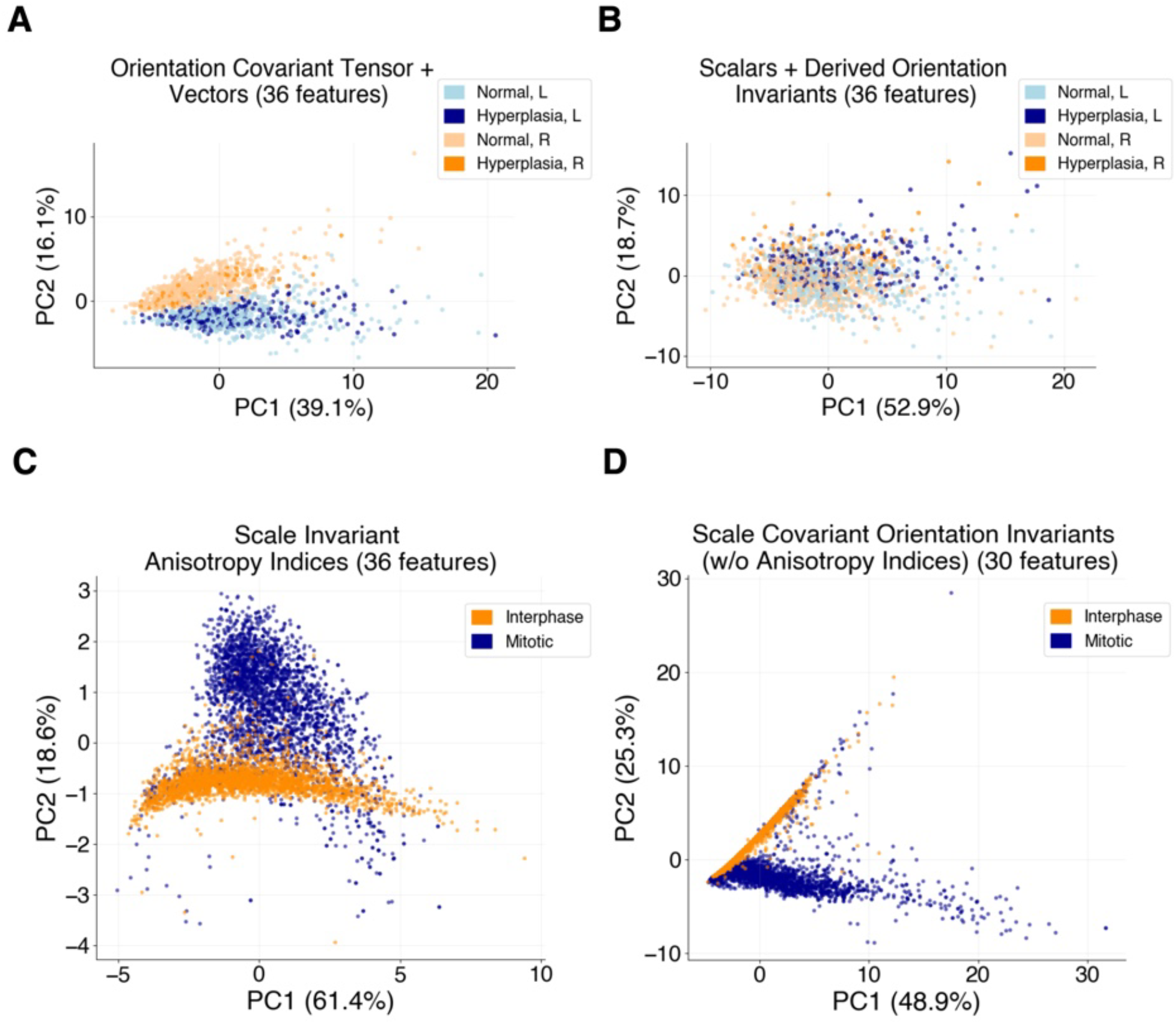
Covariant and invariant Minkowski descriptors separate biologically meaningful morphological axes. (**A, B**) PCA of Adrenal3D features. (**A**) Projection of covariant tensor and vector components resolves left (L, blue) and right (R, orange) adrenal glands into distinct clusters, confirming that covariant descriptors encode bilateral handedness. (**B**) Projection of rotation-invariant measures (scalar functionals, eigenvalues, anisotropy indices) collapses the L/R separation, as expected for orientation-independent features. (**C, D**) PCA of hiPSC nuclear features. (**C**) Projection of anisotropy indices *β* separates interphase (orange) from mitotic (dark blue) nuclei, demonstrating that scale-invariant shape descriptors alone recover cell-cycle state. (**D**) Projection of size-sensitive descriptors (scalar functionals and traces) likewise separates the two states, but additionally encodes nuclear size, which differs between interphase and mitosis.

### Scale-invariant shape distinguishes cell-cycle state in nuclei

Having shown that the covariant descriptors recover orientation on the Adrenal3D data, we now turn to the complementary behavior of scale invariance, and to a second, biologically distinct dataset (Figure 1A, bottom) to show this behavior. Nuclei undergo pronounced morphological change over the cell cycle: interphase nuclei are large, smooth and almost ellipsoid, whereas during mitosis the nuclear envelope breaks down and the chromatin condenses and reorganizes, so that shape differs markedly between interphase and mitotic states (2). This provides a natural setting to test whether Minkowski descriptors capture such biologically meaningful shape differences.

We first ask whether cell-cycle state is reflected in shape alone, independent of nuclear size. The anisotropy index *β* is scale-invariant by construction (Section 1), so any separation it produces reflects shape rather than size. Projecting the *β* values of all nuclei onto their first two principal components resolves the data into two distinct clusters, corresponding to interphase and mitotic nuclei (Fig. 2A). This demonstrates that a purely scale-invariant shape descriptor recovers cell-cycle state. The finer mitotic sub-stages do not separate cleanly, consistent with their more subtle morphological differences; the dominant, robust distinction is between interphase and mitosis.

The scalar functionals and the other rotation-invariant quantities (eigenvalues and traces) also separate interphase from mitotic nuclei. Unlike *β*, however, these quantities are not scale-invariant: they additionally encode nuclear size, which itself differs between the two states. Their separation therefore reflects a combination of size and shape, whereas the *β* result isolates the shape contribution alone. Rather than being redundant, these descriptors capture complementary aspects of morphology—rotationally-invariant shape, overall size, and topology each of which partially distinguishes the biological classes.

This multiplicity of partial, non-redundant signals is precisely what makes visual inspection insufficient: no single descriptor is evidently optimal, and the most informative combination cannot be read off by eye. We therefore turn to a supervised classification task, with a controlled ablation over descriptor subgroups, to quantify which Minkowski features carry discriminative signal and how they combine.

### MT outperforms classical and deep-learning baselines on MedMNIST classification

Minkowski tensors were extracted from triangulated surface meshes of binary segmentation masks and evaluated across the feature subgroups described in Methods. We first evaluated MT classification performance on the Adrenal3D dataset, where MT outperformed all competing feature sets—including the best convolutional baseline, exceeding the ACS ResNet by roughly nine balanced-accuracy points (0.833 vs. 0.740; Table 2) despite using a compact set of interpretable features rather than learned volumetric filters. The strongest MT subset here combined all rank-0–2 tensors with their traces and eigenvalues, the quantities that encode absolute size and elongation— appropriate for a task whose discriminative signal is glandular enlargement. To identify the most informative descriptors, we performed a permutation-importance analysis in which each feature was randomly shuffled and the resulting drop in balanced accuracy recorded, larger drops indicating greater importance. The analysis ranked the volume functional *W*_0_ highest, together with the strongly collinear trace of the solid moment tensor 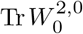, which captures the volume distribution (Figure 3A). Clinically, hyperplasia and hypertrophy of adrenal cells enlarge the gland (22), providing a volumetric diagnostic cue. Consistent with this, the trace distribution is significantly higher in hyperplastic glands, albeit with a modest effect size (Mann–Whitney *p<* 10^−12^; | Cliff’s *δ*| = 0.25, small; Figure 3B), indicating that MT recovers a biologically and clinically meaningful morphological signal and opening an avenue for interpretable, feature-level 3D morphological analysis.

**Table 2.**
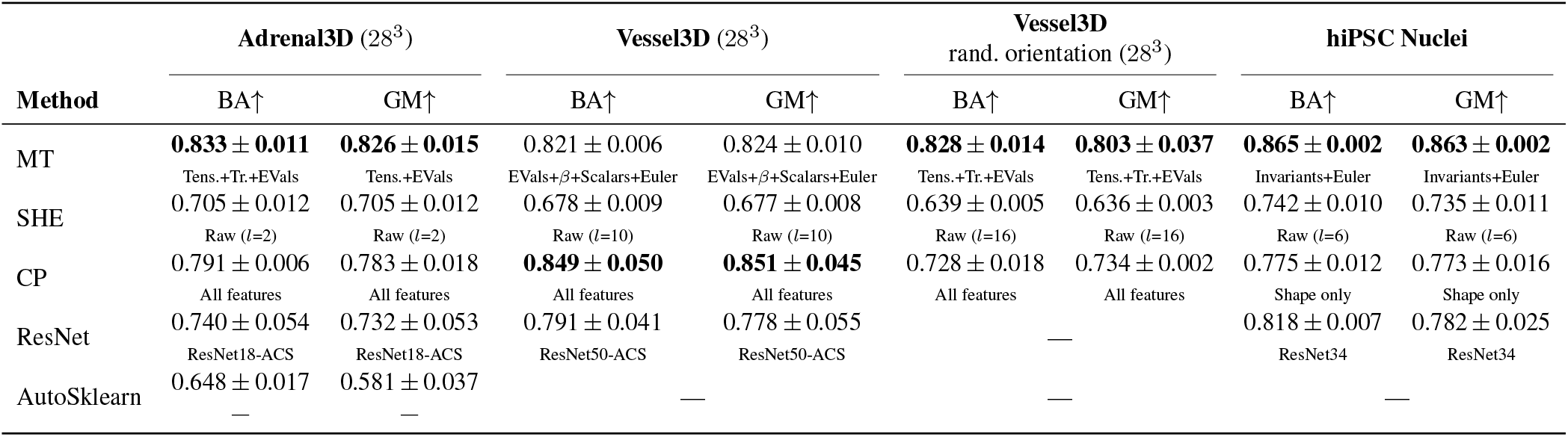
Best SVM classification performance (balanced accuracy, BA; geometric mean, GM) per method and dataset. MedMNIST datasets are evaluated at 28^3^ voxel resolution; hiPSC nuclei are at their native resolution (approximately 96 × 256 × 256; see Methods). Each cell shows mean *±* std across three independent runs and the best-performing feature subset (row labels: MT = Minkowski Tensors, SHE = spherical harmonics expansion, CP = CellProfiler, ACS = axial-coronal-sagittal convolutions (21)). **Bold** marks the highest value in each column. “—” indicates the method was not evaluated on that dataset. Vessel3D randomized orientation uses the same images as Vessel3D with a random SO(3) rotation applied independently to each object to probe rotation invariance.

**Fig. 3.**
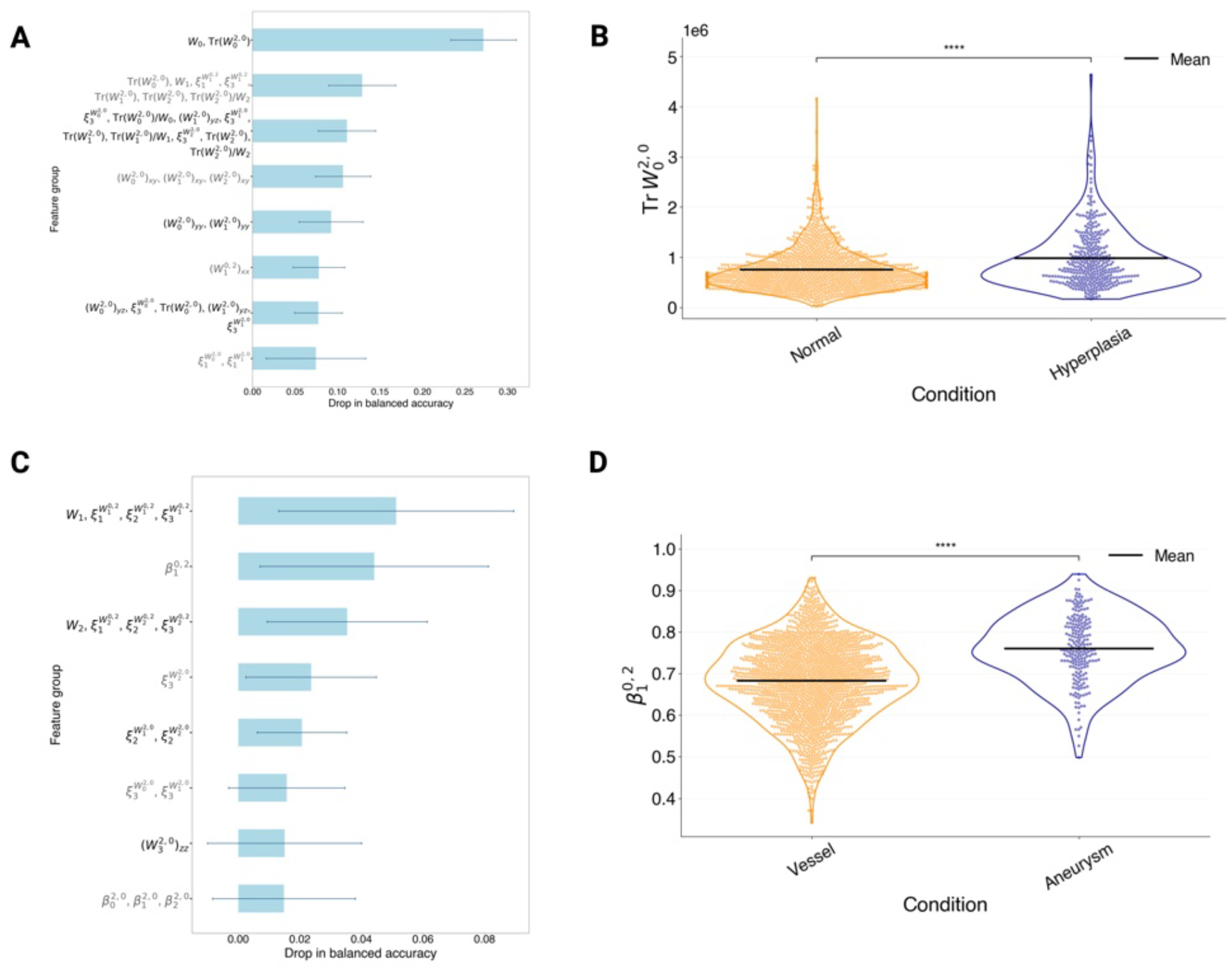
Permutation feature importance and discriminative descriptor distributions for MedMNIST datasets. (**A**) Grouped permutation importance for the Adrenal3D dataset (best MT subset: Tensors + Traces + Eigenvalues; 84 features, 56 collinear groups). Feature groups are ranked by mean drop in balanced accuracy; bars show mean ± SD across permutation repeats. The volume functional *W*_0_ and the trace 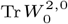 are the leading discriminative features. (**B**) Distribution of 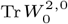 across Normal and Hyperplasia adrenal gland classes; horizontal lines indicate class means. The trace is significantly higher in hyperplastic glands (Mann–Whitney *p <* 10^−12^; Cliff’s |*δ*| = 0.25, small), consistent with glandular enlargement in hyperplasia. (**C**) Grouped permutation importance for the Vessel3D dataset (best MT subset: Eigenvalues + *β* + Scalars + Euler; 37 features, 26 collinear groups). The surface-area measure *W*_1_ and eigenvalues of 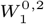, together with the anisotropy index 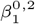, are the leading discriminative features. (**D**) Distribution of the normal distribution tensor anisotropy index 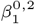 across Vessel and Aneurysm classes; horizontal lines indicate class means. The index is significantly elevated in aneurysms (Mann–Whitney *p <* 10^−18^; Cliff’s δ = −0.37, medium; Cohen’s *d* = −0.67), consistent with the more isotropic normal distribution (approaching spherical) of aneurysmal bulges relative to the elongated, near-cylindrical healthy vessels. Statistical significance: *\*\*\*\* p <* 10^−4^.

Next, we benchmarked MT and competing feature sets on the Vessel3D subset. MT achieved a balanced accuracy of 0.821 *±* 0.006, while CellProfiler outperformed MT on this dataset (0.849 ± 0.050; Table 2). Grouped permutation importance on the best-performing MT subgroup (eigenvalues, anisotropy indices, scalars, and Euler characteristic) identified the surface-area measure *W*_1_ and eigenvalues of the normal distribution tensor 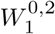 as the leading discriminative cluster, followed by the anisotropy index 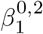 (Figure 3C). The anisotropy index 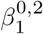 of the normal distribution tensor 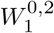 was significantly elevated in aneurysms (Mann–Whitney *p <* 10^−18^; Cliff’s *δ* = −0.37, medium; Cohen’s *d* = −0.67; Figure 3D): because 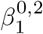 approaches its isotropic maximum when surface normals are uniformly distributed (as on a sphere) and decreases for elongated shapes, this shift is consistent with the morphological contrast between healthy, near-cylindrical vessels and the rounded, sphere-like bulging of aneurysms. Permutation importance for CellProfiler features revealed that bounding-box extent features rank among the top CP discriminators in standard orientation but are absent from the top-ranked features after random reorientation (Supplementary Fig. S1), indicating that CP’s advantage is driven by orientation-dependent position features rather than intrinsic shape. To confirm this, we created the randomized-orientation Vessel3D variant; after reorientation, MT became the top performer (0.828 ± 0.014), surpassing CP (0.728 ± 0.018; Table 2), demonstrating that MT’s discrimination reflects intrinsic geometric properties rather than dataset-specific orientation biases.

Across both datasets, feature extraction for MT and SHE requires only CPU resources and completes in seconds per sample, whereas the ResNet baselines require dedicated GPU hardware and substantially longer training times (quantitative comparison in Supplementary Table S4). Together, these results demonstrate that MT delivers robust, orientation-invariant morphological discrimination driven by biologically meaningful geometric signals, while remaining computationally accessible without specialist hardware.

### MT classifies multibody, topologically complex nuclei across the cell cycle

We next asked whether MT extends to more complex, multibody, non-zero genus morphologies that are inadequately captured by classical geometric feature sets, using high-resolution hiPSC nuclear masks spanning six cell-cycle stages (Figure 1A) (2). The nuclear masks exhibit pronounced structural variability across the cell cycle, which drives strong classification performance: MT consistently outperforms all other feature sets (Table 2), surpassing not only the classical descriptors but also the strongest deep-learning baseline, exceeding the best 3D ResNet by roughly five balanced-accuracy points and eight in geometric mean (0.865 vs. 0.818 BA, 0.863 vs. 0.782 GM; Table 2). The winning subset combined rotation invariants with the Euler characteristic—a pairing that matches the problem, since the cell-cycle stages differ chiefly in connectivity and genus, which the Euler characteristic counts directly, while the in-variant tensor quantities capture curvature without orientation dependence. The strong performance of this compact, interpretable descriptor set relative to a learned volumetric model on the most topologically complex dataset underscores the value of encoding topological and curvature features explicitly rather than relying on learned representations.

Permutation-importance analysis highlighted the rank-one curvature descriptors—in particular the eigenvalues of 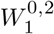—as the leading contributors to model performance (Figure 4A). Examining the largest eigenvalue 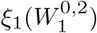 across stages, we found substantial, stage-dependent variation: the omnibus Kruskal–Wallis test was highly significant (*p <* 10^−16^), and 14 of 15 Holm-corrected pairwise Dunn comparisons reached significance, with interphase strongly separated from the mitotic stages (Cliff’s |*δ*| *>* 0.7 for most contrasts; Figure 4B). The full pairwise statistics are given in Supplementary Table S3. Because the class sizes are large and imbalanced, most pairwise comparisons reach significance; we therefore rely on Cliff’s *δ* to gauge which differences are substantial. To our knowledge, this link between MT curvature descriptors and cell-cycle stage has not previously been reported.

**Fig. 4.**
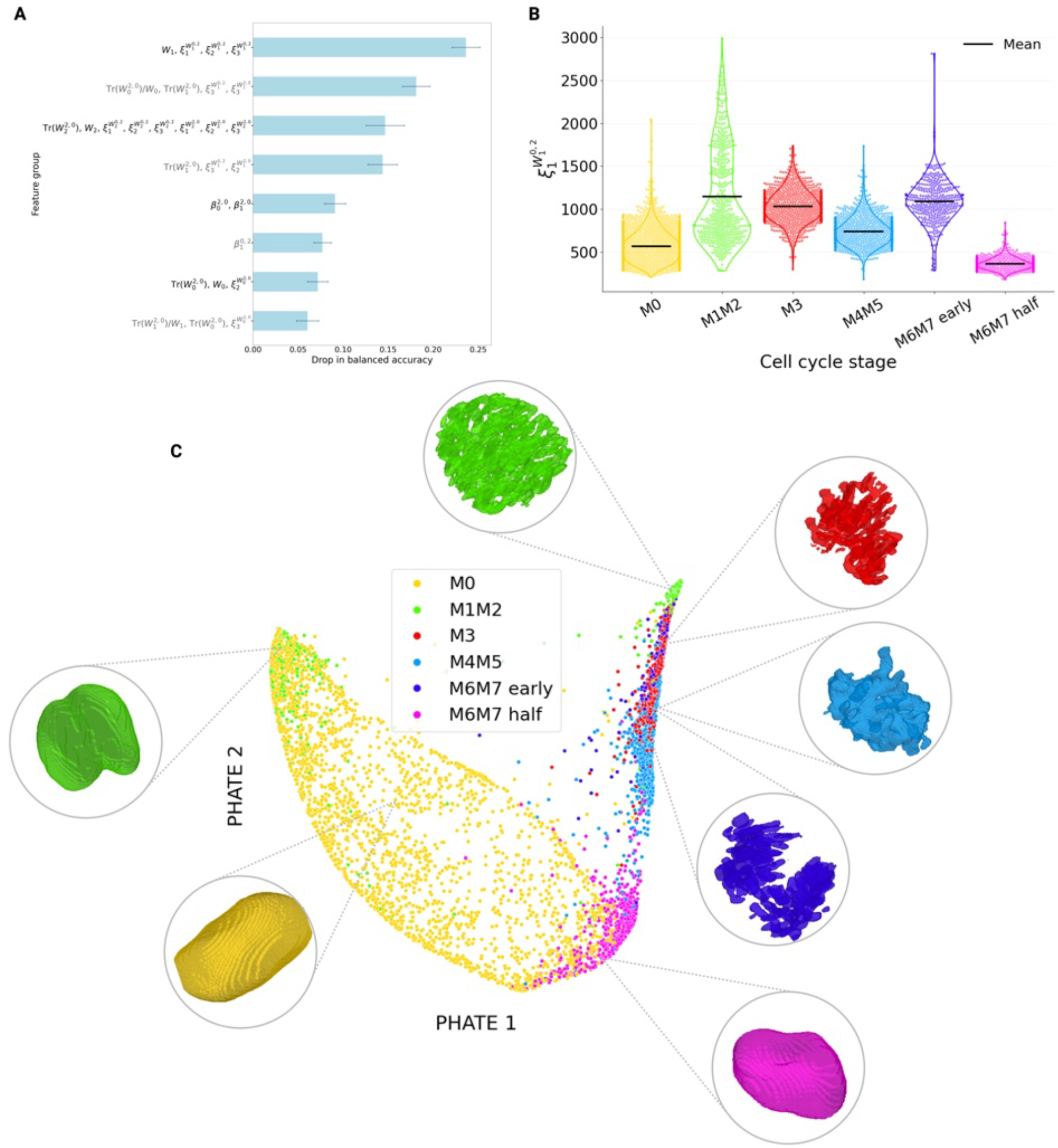
Minkowski tensor analysis of hiPSC nuclear morphology across the cell cycle. (**A**) Grouped permutation importance for the hiPSC nucleus classification task (best MT subset: Invariants + Euler; 45 features, 27 collinear groups). Feature groups are ranked by mean drop in balanced accuracy; bars show mean ± SD. Rotation-invariant curvature descriptors form the leading discriminative cluster. (**B**) Distribution of the largest eigenvalue 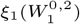 of the normal-distribution tensor 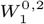 across six cell-cycle stages; horizontal lines indicate stage means. The omnibus Kruskal–Wallis test was highly significant (*p <* 10−^16^); 14 of 15 Holm-corrected pairwise Dunn comparisons reached significance, with interphase strongly separated from all mitotic stages (Cliff’s |δ| > 0.7 for most contrasts). (**C**) PHATE embedding of Minkowski features colored by cell-cycle stage, with illustrative examples of various classes as meshes. The six stages form a coherent, nearly cyclic trajectory in which adjacent stages are placed in proximity, reflecting the continuity of cell-cycle progression.

We further embedded the MT features with PHATE, a nonlinear dimensionality-reduction method that preserves local and global structure (20). MT features organized the six stages into a coherent, nearly cyclic trajectory, with adjacent stages placed close together (Figure 4C). Equivalent embeddings for the baseline descriptors are provided in Supplementary Fig. S2: CP features likewise recover a cyclic arc organization, whereas SHE embeddings yield a markedly less structured distribution with stages not clearly resolved. Together with the classification results, these findings show that MT captures topological and curvature-based complexity while exposing biologically meaningful cell-cycle dynamics.

## Discussion

In this work, we introduce and benchmark MT, a compact and interpretable feature set designed to classify 3D biological shapes across multiple scales. MT remains covariant under geometric transformations and effectively captures multi-body and topologically complex geometries. MT also does not require canonical orientation and remains robust under arbitrary rotations even without alignment preprocessing, making it well suited to morphology analysis in systems lacking a well-defined reference frame. Owing to these properties, MT achieves competitive or superior classification performance relative to existing classical and data-driven feature sets while maintaining biological interpretability. This interpretability allows MT to bridge the gap between geometric quantification and mechanistic understanding, facilitating future studies of 3D cellular and tissue morphologies. A further practical advantage is computational efficiency: MT and SHE feature extraction runs on CPU in seconds per object, whereas CellProfiler incurs greater overhead at scale and deep-learning baselines require dedicated GPU hardware and substantially longer training times (Supplementary Table S4).

Beyond overall classification performance, the permutation results and interpretability studies highlight which components of MT carry the discriminative signal. Including tensor eigenvalues, anisotropy indices, traces and trace ratios substantially improved discriminative performance, indicating that directional anisotropy and geometric variance provide complementary information beyond the base Minkowski tensors. Here, MT features are combined via simple concatenation to preserve interpretability and physical meaning; more structured or learned combinations of scalar, vectorial, and tensorial components may further improve performance.

Although powerful, MT has limitations that motivate further development. The descriptors are computed from trian-gulated surface meshes derived from binary segmentations; mesh quality and topology therefore depend directly on segmentation fidelity, and errors or artifacts in the upstream segmentation propagate into the feature values. Most Minkowski measures are not scale-invariant—only the anisotropy index β is dimensionless—so cross-dataset comparisons require careful normalization when objects differ substantially in absolute size, and incorporating additional dimensionless Minkowski combinations could better decouple shape from size, improving cross-scale generalization. The additivity property holds strictly for disjoint (non-overlapping) bodies; touching or fused segments require explicit handling of the intersection term. Finally, on strongly imbalanced datasets with rare classes, even a compact feature set may lack sufficient discriminative signal, as observed for the less-populated mitotic sub-stages.

Beyond these limitations, several directions could extend this work. Applying MT to convex hull representations would yield global morphological signatures complementary to the surface-level descriptors computed here. Leveraging the additivity property in perturbation experiments—such as drug treatment or mechanical stress assays where individual cells in a population respond variably—offers a principled way to represent the collective response as a sum of constituent contributions, capturing both aggregate and heterogeneous effects. The orientation-sensitive covariant tensor components additionally provide direct access to directional morphological information, which may be particularly valuable in developmental studies where systematic changes in cellular orientation encode spatial patterning and tissue organization; covariant descriptors could reveal such orientation-dependent dynamics without requiring explicit orientation annotation. MT may also serve as interpretable shape priors within conditional generative models, bridging the discriminative framework developed here with shape synthesis and interpolation. Application to other volumetric imaging modalities, including cryo-electron tomography and light-sheet fluorescence microscopy, represents a natural extension of the present benchmarks.

## Conclusion

We have introduced Minkowski Tensor–based feature set for classification of 3D biological morphology. Through comprehensive benchmarking across standardized and biologically complex datasets, we demonstrate that Minkowski features achieve robust, interpretable, and transformation-covariant discrimination of shape. This work establishes Minkowski features as a general framework for morphological classification and provides a simple, reproducible framework that links geometric structure to biological phenotype, empowering biologists to quantitatively compare complex shapes across scales and systems.

## Data Availability

The data underlying this article are available in the repositories listed in the Availability and Implementation section.

## Author Contributions Statement

K.I. conceived and supervised the project and acquired funding. Y.K. and K.I. designed the methodology. Y.K. conducted the experiments and analysed the results. Y.K. and K.I. wrote and reviewed the manuscript.

## ACKNOWLEDGEMENTS

The authors thank members of the Ishihara lab for helpful discussions.

## AI Disclosure

Claude Code (claude-sonnet-4-6, Anthropic) was used in the preparation of this work for the following purposes: consolidating and cleaning the original analysis codebase; completing the implementation of the 3D ResNet nuclei baseline; editing and revising manuscript text; and generating statistical tests comparing feature distributions for hiPSC data. All AI-assisted contributions were reviewed and approved by the human authors before inclusion.

## Bibliography

1. Isabel Urbina-Barreto, Frédéric Chiroleu, Romain Pinel, Louis Fréchon, Vincent Mahamadaly, Simon Elise, Michel Kulbicki, Jean-Pascal Quod, Eric Dutrieux, Rémi Garnier, J. Henrich Bruggemann, Lucie Penin, and Mehdi Adjeroh. Quantifying the shelter capacity of coral reefs using photogrammetric 3D modeling: From colonies to reefscapes. Ecological Indicators, 121:107151, 2021. doi: 10.1016/j.ecolind.2020.107151.

2. Matheus P. Viana, Jianxu Chen, Theo A. Knijnenburg, Ritvik Vasan, Calysta Yan, Joy E. Arakaki, Matte Bailey, Ben Berry, Antoine Borensztejn, et al. Integrated intracellular organization and its variations in human iPS cells. Nature, 613(7943):345–354, 2023. doi: 10.1038/s41586-022-05563-7.

3. Ishita Singh and Tanmay P. Lele. Nuclear morphological abnormalities in cancer – a search for unifying mechanisms. Results and Problems in Cell Differentiation, 70:443–467, 2022. doi: 10.1007/978-3-031-06573-6_16.

4. D’Arcy Wentworth Thompson. On Growth and Form. Cambridge University Press, Cambridge, 1997.

5. Sihong Chen, Kai Ma, and Yefeng Zheng. Med3D: Transfer learning for 3D medical image analysis, 2019. arXiv:1904.00625.

6. Xiongtao Ruan and Robert F. Murphy. Evaluation of methods for generative modeling of cell and nuclear shape. Bioinformatics, 35(14):2475–2485, 2019. doi: 10.1093/bioinformatics/bty983.

7. Anna Medyukhina, Marco Blickensdorf, Zoltán Cseresnyés, Nora Ruef, Jens V. Stein, and Marc Thilo Figge. Dynamic spherical harmonics approach for shape classification of migrating cells. Scientific Reports, 10(1), 2020. doi: 10.1038/s41598-020-62997-7.

8. Anne E. Carpenter, Thouis R. Jones, Michael R. Lamprecht, Colin Clarke, In Han Kang, Ola Friman, David A. Guertin, Joo Han Chang, Robert A. Lindquist, Jason Moffat, Polina Golland, and David M. Sabatini. CellProfiler: image analysis software for identifying and quantifying cell phenotypes. Genome Biology, 7(10):R100, 2006. doi: 10.1186/gb-2006-7-10-r100.

9. Shira Faigenbaum-Golovin and Ingrid Daubechies. Studying Morphological Variation: Exploring the Shape Space in Evolutionary Anthropology, 2024. arXiv:2410.20040.

10. Robert Osada, Thomas Funkhouser, Bernard Chazelle, and David Dobkin. Shape distributions. ACM Transactions on Graphics, 21(4):807–832, 2002. doi: 10.1145/571647.571648.

11. Claus Beisbart, Robert Dahlke, Klaus Mecke, and Herbert Wagner. Vector- and tensor-valued descriptors for spatial patterns. In Morphology of Condensed Matter, volume 600 of Lecture Notes in Physics, pages 238–260. Springer Berlin Heidelberg, 2002. doi: 10.1007/3-540-45782-8_10.

12. C. Beisbart, M. S. Barbosa, H. Wagner, and L. da F. Costa. Extended morphometric analysis of neuronal cells with Minkowski valuations. The European Physical Journal B, 52(4):531–546, 2006. doi: 10.1140/epjb/e2006-00328-1.

13. Lea Happel, Griseldis Oberschelp, Valeriia Grudtsyna, Harish P. Jain, Rastko Sknepnek, Amin Doostmohammadi, and Axel Voigt. Quantifying the shape of cells, from Minkowski tensors to p-atic orders. eLife, 14, 2025. doi: 10.7554/elife.105680.

14. Sebastien J. P. Callens, Daniel C. Tourolle née Betts, Ralph Müller, and Amir A. Zadpoor. The local and global geometry of trabecular bone. Acta Biomaterialia, 130:343–361, 2021. doi: 10.1016/j.actbio.2021.06.013.

15. Jiancheng Yang, Rui Shi, Donglai Wei, Zequan Liu, Lin Zhao, Bilian Ke, Hanspeter Pfister, and Bingbing Ni. MedMNIST v2 - a large-scale lightweight benchmark for 2d and 3d biomedical image classification. Scientific Data, 10(1):41, 2023. doi: 10.1038/s41597-022-01721-8.

16. Yajushi Khurana and Keisuke Ishihara. pykarambola: Minkowski tensor morphometry of 3d structures. bioRxiv, 2026. doi: 10.64898/2026.06.16.730752.

17. F. Pedregosa, G. Varoquaux, A. Gramfort, V. Michel, B. Thirion, O. Grisel, M. Blondel, P. Prettenhofer, R. Weiss, V. Dubourg, J. Vanderplas, A. Passos, D. Cournapeau, M. Brucher, M. Perrot, and E. Duchesnay. Scikit-learn: Machine learning in Python. Journal of Machine Learning Research, 12:2825–2830, 2011.

18. Guolin Ke, Qi Meng, Thomas Finley, Taifeng Wang, Wei Chen, Weidong Ma, Qiwei Ye, and Tie-Yan Liu. LightGBM: A highly efficient gradient boosting decision tree. In Advances in Neural Information Processing Systems, volume 30, pages 3146–3154, 2017.

19. Leo Breiman. Random forests. Machine Learning, 45(1):5–32, 2001.

20. Kevin R. Moon, David van Dijk, Zheng Wang, Scott Gigante, Daniel B. Burkhardt, William S. Chen, Kristina Yim, Antonia van den Elzen, Matthew J. Hirn, Ronald R. Coifman, Natalia B. Ivanova, Guy Wolf, and Smita Krishnaswamy. Visualizing structure and transitions in highdimensional biological data. Nature Biotechnology, 37(12):1482–1492, 2019. doi: 10.1038/s41587-019-0336-3.

21. Jiancheng Yang, Xiaoyang Huang, Yi He, Jingwei Xu, Canqian Yang, Guozheng Xu, and Bingbing Ni. Reinventing 2D convolutions for 3D images. IEEE Journal of Biomedical and Health Informatics, 25(8):3009–3018, 2021.

22. Yvonne M. Ulrich-Lai, Helmer F. Figueiredo, Michelle M. Ostrander, Dennis C. Choi, William C. Engeland, and James P. Herman. Chronic stress induces adrenal hyperplasia and hypertrophy in a subregion-specific manner. American Journal of Physiology – Endocrinology and Metabolism, 291(5):E965–E973, 2006. doi: 10.1152/ajpendo.00070.2006.

23. G. E. Schröder-Turk, W. Mickel, S. C. Kapfer, F. M. Schaller, B. Breidenbach, D. Hug, and K. Mecke. Minkowski tensors of anisotropic spatial structure. New Journal of Physics, 15 (8):083028, 2013. doi: 10.1088/1367-2630/15/8/083028.

24. Tim Head, Manoj Kumar, Holger Nahrstaedt, Gilles Louppe, and Iaroslav Shcherbatyi. scikit-optimize/scikit-optimize. 10.5281/zenodo.5565057, 2021.

25. Guillaume Lemaître, Fernando Nogueira, and Christos K. Aridas. Imbalanced-learn: A python toolbox to tackle the curse of imbalanced datasets in machine learning. Journal of Machine Learning Research, 18(17):1–5, 2017.

26. Kaiming He, Xiangyu Zhang, Shaoqing Ren, and Jian Sun. Deep residual learning for image recognition. In Proceedings of the IEEE Conference on Computer Vision and Pattern Recognition (CVPR), pages 770–778, 2016.

27. Matthias Feurer, Aaron Klein, Katharina Eggensperger, Jost Springenberg, Manuel Blum, and Frank Hutter. Efficient and robust automated machine learning. In Advances in Neural Information Processing Systems, volume 28, pages 2962–2970, 2015.

28. Henry B. Mann and Donald R. Whitney. On a test of whether one of two random variables is stochastically larger than the other. The Annals of Mathematical Statistics, 18(1):50–60, 1947.

29. Bernard L. Welch. The generalization of ‘student’s’ problem when several different population variances are involved. Biometrika, 34(1–2):28–35, 1947.

30. Frank J. Massey. The kolmogorov–smirnov test for goodness of fit. Journal of the American Statistical Association, 46(253):68–78, 1951.

31. William H. Kruskal and W. Allen Wallis. Use of ranks in one-criterion variance analysis. Journal of the American Statistical Association, 47(260):583–621, 1952.

32. Olive Jean Dunn. Multiple comparisons using rank sums. Technometrics, 6(3):241–252, 1964.

33. Sture Holm. A simple sequentially rejective multiple test procedure. Scandinavian Journal of Statistics, 6(2):65–70, 1979.

34. Norman Cliff. Dominance statistics: Ordinal analyses to answer ordinal questions. Psychological Bulletin, 114(3):494–509, 1993.

35. Jacob Cohen. Statistical Power Analysis for the Behavioral Sciences. Lawrence Erlbaum Associates, Hillsdale, NJ, 2nd edition, 1988.

36. Pauli Virtanen, Ralf Gommers, Travis E. Oliphant, Matt Haberland, Tyler Reddy, David Cournapeau, Evgeni Burovski, Pearu Peterson, Warren Weckesser, Jonathan Bright, Stéfan J. van der Walt, Matthew Brett, Joshua Wilson, K. Jarrod Millman, Nikolay Mayorov, Andrew R. J. Nelson, Eric Jones, Robert Kern, Eric Larson, C. J. Carey, ?Ilhan Polat, Yu Feng, Eric W. Moore, Jake VanderPlas, Denis Laxalde, Josef Perktold, Robert Cimrman, Ian Henriksen, E. A. Quintero, Charles R. Harris, Anne M. Archibald, Antônio H. Ribeiro, Fabian Pedregosa, Paul van Mulbregt, and SciPy 1.0 Contributors. SciPy 1.0: Fundamental algorithms for scientific computing in Python. Nature Methods, 17(3):261–272, 2020. doi: 10.1038/s41592-019-0686-2.

