## Supplementary File for "Minkowski Tensors as a Lightweight and Interpretable Representation for Three-Dimensional Morphology"

### Supplementary Figures and Tables

#### Supplementary Methods.

##### *Datasets: Additional Details.*

**MedMNIST.** MedMNIST provides preprocessed and segmented 3D medical imaging datasets with predefined training, validation, and test splits, enabling reproducible benchmarking across studies (15). Each dataset contains approximately $10^3$ volumetric samples distributed across two imbalanced classes (the normal or healthy class predominating) and follows a standardized 7:2:1 split for training, testing, and validation. The datasets are provided at two spatial resolutions ($28^3$ and $64^3$), allowing assessment of feature robustness across voxel scales.

**Allen Cell Institute hiPSC Dataset.** The dataset consists of high-resolution fluorescence microscopy images of human iPS cells with nuclear and cellular segmentations annotated for different stages of the cell cycle (2); we used only the nuclear segmentation masks. The dataset comprises 5606 segmented nuclei distributed across six cell-cycle classes — Interphase, Early Prometaphase, Prometaphase/Metaphase, Prophase, and two Anaphase/Telophase/Cytokinesis stages (half and early) — forming a multiclass classification task with Interphase predominating. We used isotropic nuclear masks of varying spatial extent (approximately $300^3$) and standardized them to a fixed array size (see Preprocessing below).

***Preprocessing: Additional Details.*** Minkowski tensors are translation-covariant or invariant, so their values depend on the choice of coordinate origin. We use pykarambola (16), which resolves this by computing tensors about the mesh centroid by default. To give every descriptor method in our benchmark the same origin convention, we translated each mask so that its center of mass coincided with the array center; this was a pure translation with no rotation or axis alignment, so each object's native orientation is preserved. For the MedMNIST subsets the center-of-mass offsets were below two voxels, making this step nearly an identity there; we applied it uniformly nonetheless for consistency across datasets.

The hiPSC nuclei required an additional standardization step. The masks were obtained already resampled to isotropic voxels (2), but individual nuclear crops varied widely in extent. Because our deep-learning baselines consume the binary volume directly, they require fixed-size input images; we standardized every nucleus to a common array size to prevent voxel-based models from keying on crop or bounding-box dimensions rather than on intrinsic nuclear geometry. We followed the downsampling and size-standardization scheme used by Viana et al. (2) with a few modifications: each mask was first downsampled by a factor of $0.5$ using nearest-neighbor interpolation (preserving strict binary values), bringing per-axis extents to roughly 41–108 (Z), 100–307 (Y), and 105–310 (X) voxels. We then center-padded each volume with background to a fixed $96 \times 256 \times 256$ ($Z \times Y \times X$) array — center-cropping the largest $\sim 2\%$ of volumes — with the target chosen to enclose $98\%$ of nuclei without truncation. We padded rather than rescaled each nucleus to preserve absolute size and aspect ratio, which change meaningfully across the cell cycle and feed directly into size-sensitive geometric descriptors; rescaling to a cube would erase that information. Because padding adds only background voxels, it leaves Minkowski tensors and other foreground-based descriptors unchanged.

***Minkowski Tensor Mathematical Definitions.*** While Minkowski tensors are defined at arbitrary rank, we restrict the present work to ranks 0 through 2 for three reasons: tensors of higher rank become increasingly sensitive to discretization noise on triangulated surfaces; ranks 0–2 admit direct geometric interpretation (object size, orientation, and anisotropy) whereas higher-rank components lack comparably intuitive meaning; and restricting to low ranks keeps the Cartesian representation compact and non-redundant. In three dimensions these comprise four rank-0 scalars (the Minkowski functionals), four rank-1 vectors, and six linearly independent rank-2 symmetric tensors, denoted $W_\nu^{r,s}$, where $\nu$ specifies the geometric measure type ($\nu = 0$: volume, $\nu = 1$: surface area, $\nu = 2$: mean curvature, $\nu = 3$: Gaussian curvature weighted) and the total tensor rank is $r + s$; integral definitions are given in Table S1. In practice, for volumetric data these quantities are computed in discrete form on triangulated surface meshes.

For each rank-2 Minkowski tensor the anisotropy index is

$$\beta_\nu^{r,s} = \left| \frac{\xi_1}{\xi_3} \right| \in [0, 1],$$

where $|\xi_1| \leq |\xi_2| \leq |\xi_3|$ are the tensor eigenvalues; a value of unity indicates perfect isotropy and smaller values indicate anisotropy. The trace is $\mathrm{Tr}(W) = \sum_{i=1}^{3} W_{ii} = \xi_1 + \xi_2 + \xi_3$ and the determinant is $\det(W) = \xi_1 \xi_2 \xi_3$; both are rotationally invariant. For position-type rank-2 tensors ($W_\nu^{2,0}$), we additionally compute the trace ratio $\mathrm{Tr}(W_\nu^{2,0})/W_\nu$, a rotationally invariant measure of how tensorial content is concentrated relative to the isotropic scalar magnitude; this ratio is not defined for normal-type tensors $W_\nu^{0,2}$ whose trace is identically equal to the corresponding scalar. Beyond individual measures, Minkowski tensors as a family are additive:

$$W_\nu^{r,s}(K \cup K') = W_\nu^{r,s}(K) + W_\nu^{r,s}(K') - W_\nu^{r,s}(K \cap K'),$$

for convex $K$, $K'$ (23).

| | |
|---|---|
| Scalar measures (rank zero) | |
| $W_0$ | $\int_K dV$ |
| $W_1$ | $\frac{1}{3}\int_{\partial K} dA$ |
| $W_2$ | $\frac{1}{3}\int_{\partial K} H\, dA$ |
| $W_3$ | $\frac{1}{3}\int_{\partial K} K_G\, dA = \frac{2\pi}{3}\chi$ |
| Vectorial measures (rank one) | |
| $(W_0^{1,0})_i$ | $\int_K \mathbf{x}_i dV$ |
| $(W_1^{1,0})_i$ | $\frac{1}{3}\int_{\partial K} \mathbf{x}_i dA$ |
| $(W_2^{1,0})_i$ | $\frac{1}{3}\int_{\partial K} H\, \mathbf{x}_i dA$ |
| $(W_3^{1,0})_i$ | $\frac{1}{3}\int_{\partial K} K_G\, \mathbf{x}_i dA$ |
| Tensorial measures (rank two) | |
| $(W_0^{2,0})_{ij}$ | $\int_K \mathbf{x}_i \mathbf{x}_j dV$ |
| $(W_1^{2,0})_{ij}$ | $\frac{1}{3}\int_{\partial K} \mathbf{x}_i \mathbf{x}_j dA$ |
| $(W_2^{2,0})_{ij}$ | $\frac{1}{3}\int_{\partial K} H\, \mathbf{x}_i \mathbf{x}_j dA$ |
| $(W_3^{2,0})_{ij}$ | $\frac{1}{3}\int_{\partial K} K_G\, \mathbf{x}_i \mathbf{x}_j dA$ |
| $(W_1^{0,2})_{ij}$ | $\frac{1}{3}\int_{\partial K} \mathbf{n}_i \mathbf{n}_j dA$ |
| $(W_2^{0,2})_{ij}$ | $\frac{1}{3}\int_{\partial K} H\, \mathbf{n}_i \mathbf{n}_j dA$ |

**Table S1.** Minkowski functionals (MF) and Minkowski tensors (MT) and their integral definitions in 3D for a body $K$ with smooth boundary $\partial K$. The mean and Gaussian curvatures on $\partial K$ are $H$ and $K_G$, respectively, and $\chi$ is the Euler characteristic. Physical interpretations and transformation properties of each quantity are given in Table 1.

***Feature Set Construction and Ablation Design.*** The Minkowski tensor computation yields, for each object, the raw scalar, vector, and rank-2 tensor components together with derived rotation-invariant quantities (eigenvalues, anisotropy indices $\beta$, traces, determinants, and trace ratios). Rather than committing to a single feature vector, we defined a family of nested and overlapping feature subgroups and evaluated each under an identical classification protocol (Section ). The purpose of this ablation is not to identify a single best-performing subset, but to attribute the discriminative signal to specific components of the descriptor and to characterize the trade-off between dimensionality and performance.
These subgroups are organized along interpretable axes: reduction to invariants, augmentation of the raw components, and topological enrichment. They are not intended as a controlled factorial over individual properties, since several invariants necessarily co-vary; the aim is to attribute discriminative signal to broad classes of feature rather than to isolate the effect of any single property.

**Raw tensorial baseline.** The reference subgroup of Tensors comprises the four scalar functionals, the four rank-1 vectors, and the six rank-2 tensors in their full matrix form. This retains all orientation-dependent (covariant) information and serves as the baseline.

**Reduction to invariant summaries.** Our first line of ablation asks whether the orientation-specific matrix components are necessary, or whether the rotation-invariant summaries of each tensor suffice. We therefore constructed subgroups built exclusively from invariant quantities: the eigenvalues alone, the anisotropy indices alone, the traces alone, traces with determinants, and their combinations. These sets are substantially lower-dimensional than the raw tensorial baseline; the most compact contains only six features, isolating the contribution of each invariant. In terms of Table 1, this axis progressively discards the covariant, scale-dependent raw components in favor of the rotationally invariant summaries (eigenvalues, traces) and the additionally scale-invariant ratios ($\beta$), testing how much discriminative signal survives when only the invariant part of each tensor is retained.

**Augmentation of the raw components.** A complementary line asks whether adding invariant summaries on top of the raw components improves discrimination beyond the baseline. The corresponding subgroups augment the raw tensors with traces alone, with eigenvalues alone, with $\beta$ alone, with traces and eigenvalues together, and with all three combined. Traces are included as an explicit augmentation because, unlike eigenvalues and $\beta$, they are additive (Table 1) and capture a distinct aspect of tensor magnitude that the full matrix components do not separately emphasize in a rotation-invariant form.

**Topological augmentation.** To assess the contribution of explicit topological information, we defined subgroups that add the Euler-characteristic–related term and associated components to the invariant sets. These subgroups isolate the effect of the $W_3$-

derived, hole-sensitive quantities identified in Table 1, which are the only components carrying explicit topological information; their inclusion tests whether topological signal contributes beyond the geometric and anisotropic features on datasets containing non-zero genus morphologies.

**Full invariant set.** Finally, we aggregate the scalars, eigenvalues, $\beta$ indices, linearly independent traces, and trace ratios into a single 36-dimensional rotation-invariant descriptor. Each subgroup was evaluated with the identical pipeline and repeated-run protocol described in Section , so that differences in performance reflect the feature content alone rather than differences in model or tuning. Full results for all subgroups and both classifiers (SVM and LightGBM) are provided in the Supplementary Data Files (`svm_all_scores.csv` and `lgbm_all_scores.csv`).

***Classification Pipeline Details.*** Each feature set is standardized and passed to an SVM classifier (17). Hyperparameters (the penalty $C$, kernel, and kernel coefficient $\gamma$) are tuned by Bayesian optimization (24), which uses a probabilistic surrogate model to search the configuration space efficiently, with 50 iterations of five-fold stratified cross-validation on the training set. For class-imbalanced datasets the SVM is wrapped in a balanced bagging ensemble that resamples each bootstrap to equalize class frequencies (25); balanced datasets use a standard bagging ensemble. To confirm that our findings reflect the feature sets rather than a particular classifier, we repeated the analysis with a LightGBM gradient-boosted tree classifier (18); LightGBM corrects for class imbalance internally through class weighting and therefore requires no balanced bagging wrapper. For each feature set and classifier, the complete procedure—hyperparameter selection, refitting on the combined training and validation data, and evaluation on the held-out test set—was repeated three times with independent random seeds, and we report the mean and standard deviation across runs.
Classification performance is evaluated using balanced accuracy (BA) and the geometric mean (G-mean), both appropriate for imbalanced data. For a problem with $N$ classes, let $\mathrm{recall}_i$ denote the recall (true positive rate) of class $i$. Balanced accuracy is the mean per-class recall,

$$\mathrm{BA} = \frac{1}{N}\sum_{i=1}^{N}\mathrm{recall}_i,$$

and the geometric mean is the $N$th root of their product,

$$\text{G-mean} = \left(\prod_{i=1}^{N}\mathrm{recall}_i\right)^{1/N}.$$

Both weight all classes equally regardless of size; G-mean is the more stringent, as it collapses to zero if any single class is never recalled.

***Benchmarking and Deep Learning Baseline Details.*** We benchmark the Minkowski feature sets against CellProfiler and spherical harmonic descriptors. CellProfiler features (8) were extracted with the `MeasureSizeShape` module and evaluated as several subgroups: the full feature set, shape features only, location features only, and individual interpretable descriptors (surface area, solidity, and their combination). Spherical harmonic descriptors were computed with the `aics-shparam` library (2); we evaluated expansions from $l_{\max} = 1$ to $l_{\max} = 16$, as classification performance plateaued at low $l_{\max}$ and declined at higher orders, consistent with high-order coefficients encoding increasingly noise-dominated detail. All descriptor sets were processed through the identical classification pipeline described in Section , so that performance differences are attributable to the feature set rather than to the classifier or its tuning.
We additionally compared against deep-learning baselines, handled differently for the two data sources. For the MedMNIST subsets (Adrenal3D, Vessel3D) we adopt the published benchmark results (15), reporting the strongest performers ResNet-18 and ResNet-50 with axial-coronal-sagittal (ACS) convolutions (21, 26) and Auto-sklearn (27). For the hiPSC nuclei, for which no established benchmark exists, we trained native 3D ResNet models, adapting the architecture of the AllenCell `image_classifier_3d` framework (2) and the ResNet implementation of the MedMNIST benchmark (15); we trained ResNet-34, ResNet-50, and ResNet-101 across three random seeds; all three are reported in Supplementary Table S6, and ResNet-34 (the strongest by balanced accuracy) is included in Table 2. All deep baselines were evaluated with the same balanced accuracy and geometric mean metrics as the Minkowski pipeline, recomputed from per-sample predictions where necessary. Two differences in protocol should be noted: the trained ResNet models are fit on the training split alone (with the validation split used only to select the best epoch), whereas the descriptor-based pipelines are refit on the combined training and validation data after hyperparameter selection; and variance for deep models reflects independent training runs while descriptor-pipeline variance reflects re-seeded hyperparameter selection and fitting.

***Permutation Importance Details.*** Permutation feature importance (19) is quantified as the decrease in balanced accuracy when a feature's values are randomly permuted while all others are held fixed, computed via five-fold stratified cross-validation with importance measured on each held-out fold and aggregated as the mean across folds; the classification pipeline is otherwise

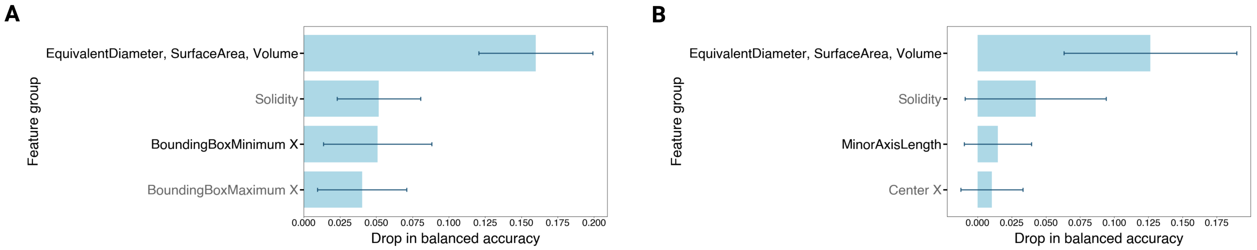


**Fig. S1.** Grouped permutation importance (top 4 features) for CP on Vessel3D in standard orientation (**A**) and after random SO(3) re-orientation (**B**). Features are ranked by mean drop in balanced accuracy; bars show mean ± SD. BoundingBoxMinimum X and BoundingBoxMaximum X rank among the top discriminative features in standard orientation but are absent from the top-4 after random reorientation, reflecting their dependence on absolute image-frame alignment rather than intrinsic object geometry.

unchanged. Because many Minkowski descriptors are strongly collinear — eigenvalues, traces, anisotropy indices, and raw components can be functions of the same underlying tensor — single-feature permutation can understate importance, as correlated features compensate for the permuted one. We therefore additionally report a grouped permutation importance in which strongly correlated features ($|r| > 0.9$) are permuted jointly, so that the reported value reflects the contribution of each correlated cluster rather than being diluted across redundant descriptors. Importance was evaluated with the same feature matrix used to train the final model (training and validation combined), with five permutation repeats per feature.

***Statistical Analysis Details.*** To probe biological interpretability, we examined the highest-ranked features from the permutation importance analysis and characterized how their distributions vary across classes. Throughout, we adopted a rank-based testing family so that results remain directly comparable between the two-class MedMNIST datasets and the six-class hiPSC nuclei dataset. All tests were two-sided with a significance threshold of $\alpha = 0.05$.

For the two-class datasets (Adrenal3D and Vessel3D) we used the Mann–Whitney $U$ test (28) as the primary assessment, since it is sensitive to a shift in location without assuming normality. This was complemented by Welch's $t$-test (29), which does not assume equal variances, and by the two-sample Kolmogorov–Smirnov test (30), which is sensitive to differences in the full shape of the distribution rather than location alone. As each feature involves a single comparison in this setting, no multiple-comparison correction was applied.

For the six-class hiPSC nuclei dataset we first applied the Kruskal–Wallis test (31) as an omnibus check for any difference among the mitotic stages. Where the omnibus test was significant, we performed Dunn's post-hoc test (32) across all $\binom{6}{2} = 15$ pairwise comparisons and controlled the family-wise error rate using the Holm–Bonferroni procedure (33); reported pairwise $p$-values are the Holm-adjusted values.

Because several of the class distributions are strongly imbalanced (the interphase population is substantially larger than the mitotic stages), statistical significance alone is a weak indicator of biological relevance: even negligible differences are readily detected at large sample sizes. We therefore report effect sizes alongside every test: Cliff's $\delta$ (34) as the primary rank-based effect-size measure (interpreted using conventional thresholds of $|\delta| < 0.147$ negligible, $< 0.33$ small, $< 0.474$ medium, and larger values large), with Cohen's $d$ (35) as a complementary standardized mean difference. For pairwise comparisons in which the adjusted $p$-value underflowed to zero owing to very large rank separations, Cliff's $\delta$ was used to order the comparisons by effect magnitude. All tests were computed using SciPy (36).

Comprehensive ablation results across all subgroups for both classifiers are provided as Supplementary Data Files: `svm_all_scores.csv` (SVM) and `lgbm_all_scores.csv` (LightGBM).

### Supplementary References


1. Isabel Urbina-Barreto, Frédéric Chiroleu, Romain Pinel, Louis Fréchon, Vincent Mahamadaly, Simon Elise, Michel Kulbicki, Jean-Pascal Quod, Eric Dutrieux, Rémi Garnier, J. Henrich Bruggemann, Lucie Penin, and Mehdi Adjeroh. Quantifying the shelter capacity of coral reefs using photogrammetric 3D modeling: From colonies to reefscapes. *Ecological Indicators*, 121:107151, 2021. doi: 10.1016/j.ecolind.2020.107151.
2. Matheus P. Viana, Jianxu Chen, Theo A. Knijnenburg, Ritvik Vasan, Calysta Yan, Joy E. Arakaki, Matte Bailey, Ben Berry, Antoine Borensztejn, et al. Integrated intracellular organization and its variations in human iPS cells. *Nature*, 613(7943):345–354, 2023. doi: 10.1038/s41586-022-05563-7.
3. Ishita Singh and Tanmay P. Lele. Nuclear morphological abnormalities in cancer – a search for unifying mechanisms. *Results and Problems in Cell Differentiation*, 70:443–467, 2022. doi: 10.1007/978-3-031-06573-6_16.
4. D'Arcy Wentworth Thompson. *On Growth and Form*. Cambridge University Press, Cambridge, 1997.
5. Sihong Chen, Kai Ma, and Yefeng Zheng. Med3D: Transfer learning for 3D medical image analysis, 2019. arXiv:1904.00625.
6. Xiongtao Ruan and Robert F. Murphy. Evaluation of methods for generative modeling of cell and nuclear shape. *Bioinformatics*, 35(14):2475–2485, 2019. doi: 10.1093/bioinformatics/bty983.
7. Anna Medyukhina, Marco Blickensdorf, Zoltán Cseresnyés, Nora Ruef, Jens V. Stein, and Marc Thilo Figge. Dynamic spherical harmonics approach for shape classification of migrating cells. *Scientific Reports*, 10(1), 2020. doi: 10.1038/s41598-020-62997-7.
8. Anne E. Carpenter, Thouis R. Jones, Michael R. Lamprecht, Colin Clarke, In Han Kang, Ola Friman, David A. Guertin, Joo Han Chang, Robert A. Lindquist, Jason Moffat, Polina Golland, and David M. Sabatini. CellProfiler: image analysis software for identifying and quantifying cell phenotypes. *Genome Biology*, 7(10):R100, 2006. doi: 10.1186/gb-2006-7-10-r100.
9. Shira Faigenbaum-Golovin and Ingrid Daubechies. Studying Morphological Variation: Exploring the Shape Space in Evolutionary Anthropology, 2024. arXiv:2410.20040.
10. Robert Osada, Thomas Funkhouser, Bernard Chazelle, and David Dobkin. Shape distributions. *ACM Transactions on Graphics*, 21(4):807–832, 2002. doi: 10.1145/571647.571648.

**Table S2.** All Minkowski tensor (MT) and CellProfiler (CP) feature subgroups evaluated in the benchmark. “Tensors” denotes all rank-0 through rank-2 Minkowski tensors: 4 scalars ($W_0$–$W_3$), 4 rank-1 vectors ($W_\nu^{1,0}$, 3 components each), and 6 rank-2 tensor matrices ($W_\nu^{r,s}$, 6 upper-triangular components each). Non-redundant traces are $\mathrm{Tr}(W_\nu^{2,0})$ for $\nu = 0,1,2,3$; $\mathrm{Tr}(W_1^{0,2})$ and $\mathrm{Tr}(W_2^{0,2})$ are omitted because they are proportional to $W_1$ and $W_2$ respectively. Trace ratios are $\mathrm{Tr}(W_\nu^{2,0})/W_\nu$ for $\nu = 0,1,2,3$. Euler family: $W_3$ scalar (1), $W_3^{1,0}$ vector components (3), $W_3^{0,2}$ matrix components (6). $N$: total number of features.

| Subgroup | Features included | $N$ |
|---|---|---|
| *MT — raw tensorial components* | | |
| Tensors | 4 scalars + 4 vectors (3 comp. each) + 6 rank-2 matrices (6 unique each) | 52 |
| Tensors+$\beta$ | Tensors + 6 anisotropy indices $\beta$ | 58 |
| Tensors+EVals | Tensors + 18 eigenvalues (3 per rank-2 tensor) | 70 |
| Tensors+EVals+$\beta$ | Tensors + 18 eigenvalues + 6 $\beta$ | 76 |
| Tensors+Traces | Tensors + 4 non-redundant traces + 4 trace ratios | 60 |
| Tensors+Traces+EVals | Tensors+Traces + 18 eigenvalues + 6 $\beta$ | 84 |
| *MT — invariant summaries only* | | |
| EVals only | 18 eigenvalues (3 per rank-2 tensor) | 18 |
| Beta only | 6 anisotropy indices $\beta$ | 6 |
| Traces only | 6 traces (one per rank-2 tensor) | 6 |
| EVals+$\beta$ | 18 eigenvalues + 6 $\beta$ | 24 |
| Traces+Det | 6 traces + 6 determinants | 12 |
| Traces+Det+$\beta$ | 6 traces + 6 determinants + 6 $\beta$ | 18 |
| Scalars+Traces | 4 scalars + 4 non-redundant traces | 8 |
| Invariants | 4 scalars + 18 eigenvalues + 6 $\beta$ + 4 non-redundant traces + 4 trace ratios | 36 |
| *MT — Euler-characteristic augmented* | | |
| EVals+$\beta$+Euler | 18 eigenvalues + 6 $\beta$ + Euler family (1+3+6 components) | 34 |
| EVals+$\beta$+Scalars+Euler | 18 eigenvalues + 6 $\beta$ + 4 scalars (incl. $W_3$) + $W_3^{1,0}$ (3) + $W_3^{0,2}$ (6) | 37 |
| Invariants+Euler | Invariants + $W_3^{1,0}$ vector (3) + $W_3^{0,2}$ raw matrix components (6) | 45 |
| Invariants+Euler$-\beta$ | Invariants (excl. $\beta$) + $W_3^{1,0}$ vector (3) + $W_3^{0,2}$ raw matrix components (6) | 39 |
| *CellProfiler* | | |
| CP All features | 18 AreaShape + 3 Location (Center X/Y/Z) + 1 object count | 22 |
| CP Shape only | 18 AreaShape features (bounding box, extent, diameter, surface area, volume, solidity) | 18 |
| CP Location only | 3 location features (Center X, Y, Z) | 3 |
| CP Surface area | Surface area only | 1 |
| CP Solidity | Solidity only | 1 |
| CP SA+Solidity | Surface area + Solidity | 2 |


11. Claus Beisbart, Robert Dahlke, Klaus Mecke, and Herbert Wagner. Vector- and tensor-valued descriptors for spatial patterns. In *Morphology of Condensed Matter*, volume 600 of *Lecture Notes in Physics*, pages 238–260. Springer Berlin Heidelberg, 2002. doi: 10.1007/3-540-45782-8_10.
12. C. Beisbart, M. S. Barbosa, H. Wagner, and L. da F. Costa. Extended morphometric analysis of neuronal cells with Minkowski valuations. *The European Physical Journal B*, 52(4):531–546, 2006. doi: 10.1140/epjb/e2006-00328-1.
13. Lea Happel, Griseldis Oberschelp, Valeriia Grudtsyna, Harish P. Jain, Rastko Sknepnek, Amin Doostmohammadi, and Axel Voigt. Quantifying the shape of cells, from Minkowski tensors to p-atic orders. *eLife*, 14, 2025. doi: 10.7554/elife.105680.
14. Sebastien J. P. Callens, Daniel C. Tourolle née Betts, Ralph Müller, and Amir A. Zadpoor. The local and global geometry of trabecular bone. *Acta Biomaterialia*, 130:343–361, 2021. doi: 10.1016/j.actbio.2021.06.013.
15. Jiancheng Yang, Rui Shi, Donglai Wei, Zequan Liu, Lin Zhao, Bilian Ke, Hanspeter Pfister, and Bingbing Ni. MedMNIST v2 - a large-scale lightweight benchmark for 2d and 3d biomedical image classification. *Scientific Data*, 10(1):41, 2023. doi: 10.1038/s41597-022-01721-8.
16. Yajushi Khurana and Keisuke Ishihara. pykarambola: Minkowski tensor morphometry of 3d structures. *bioRxiv*, 2026. doi: 10.64898/2026.06.16.730752.
17. F. Pedregosa, G. Varoquaux, A. Gramfort, V. Michel, B. Thirion, O. Grisel, M. Blondel, P. Prettenhofer, R. Weiss, V. Dubourg, J. Vanderplas, A. Passos, D. Cournapeau, M. Brucher, M. Perrot, and E. Duchesnay. Scikit-learn: Machine learning in Python. *Journal of Machine Learning Research*, 12:2825–2830, 2011.
18. Guolin Ke, Qi Meng, Thomas Finley, Taifeng Wang, Wei Chen, Weidong Ma, Qiwei Ye, and Tie-Yan Liu. LightGBM: A highly efficient gradient boosting decision tree. In *Advances in Neural Information Processing Systems*, volume 30, pages 3146–3154, 2017.
19. Leo Breiman. Random forests. *Machine Learning*, 45(1):5–32, 2001.
20. Kevin R. Moon, David van Dijk, Zheng Wang, Scott Gigante, Daniel B. Burkhardt, William S. Chen, Kristina Yim, Antonia van den Elzen, Matthew J. Hirn, Ronald R. Coifman, Natalia B. Ivanova, Guy Wolf, and Smita Krishnaswamy. Visualizing structure and transitions in high-dimensional biological data. *Nature Biotechnology*, 37(12):1482–1492, 2019. doi: 10.1038/s41587-019-0336-3.
21. Jiancheng Yang, Xiaoyang Huang, Yi He, Jingwei Xu, Canqian Yang, Guozheng Xu, and Bingbing Ni. Reinventing 2D convolutions for 3D images. *IEEE Journal of Biomedical and Health Informatics*, 25(8):3009–3018, 2021.
22. Yvonne M. Ulrich-Lai, Helmer F. Figueiredo, Michelle M. Ostrander, Dennis C. Choi, William C. Engeland, and James P. Herman. Chronic stress induces adrenal hyperplasia and hypertrophy in a subregion-specific manner. *American Journal of Physiology – Endocrinology and Metabolism*, 291(5):E965–E973, 2006. doi: 10.1152/ajpendo.00070.2006.
23. G. E. Schröder-Turk, W. Mickel, S. C. Kapfer, F. M. Schaller, B. Breidenbach, D. Hug, and K. Mecke. Minkowski tensors of anisotropic spatial structure. *New Journal of Physics*, 15(8):083028, 2013. doi: 10.1088/1367-2630/15/8/083028.
24. Tim Head, Manoj Kumar, Holger Nahrstaedt, Gilles Louppe, and Iaroslav Shcherbatyi. scikit-optimize/scikit-optimize. https://doi.org/10.5281/zenodo.5565057, 2021.
25. Guillaume Lemaître, Fernando Nogueira, and Christos K. Aridas. Imbalanced-learn: A python toolbox to tackle the curse of imbalanced datasets in machine learning. *Journal of Machine Learning*

**Table S3.** Dunn post-hoc pairwise comparison statistics for the largest eigenvalue $\xi_1(W_1^{0,2})$ across all 15 cell-cycle stage pairs in the Allen Cell hiPSC nuclei dataset. $p$-values are Holm–Bonferroni corrected; Cliff's $\delta$ magnitude thresholds: negligible $|\delta| < 0.147$, small $< 0.33$, medium $< 0.474$, large $\geq 0.474$. Values of $p_{\text{Holm}} = 0$ indicate underflow below machine precision ($< 10^{-300}$). Stage codes: M0 = Interphase; M1M2 = Prophase; M3 = Early Prometaphase; M4M5 = Prometaphase/Metaphase; M6M7e = Anaphase/Telophase/Cytokinesis Early; M6M7h = Anaphase/Telophase/Cytokinesis Half.

| Stage A | Stage B | $z$ | $p_{\text{Holm}}$ | Cliff's $\delta$ | Magnitude |
|---|---|---|---|---|---|
| M3 | M6M7h | $+44.29$ | $< 10^{-300}$ | $+0.996$ | large |
| M0 | M3 | $-38.48$ | $< 10^{-300}$ | $-0.914$ | large |
| M6M7e | M6M7h | $+36.14$ | $< 10^{-300}$ | $+0.976$ | large |
| M1M2 | M6M7h | $+34.51$ | $< 10^{-300}$ | $+0.977$ | large |
| M4M5 | M6M7h | $+29.91$ | $< 10^{-300}$ | $+0.973$ | large |
| M0 | M6M7e | $-27.77$ | $< 10^{-300}$ | $-0.901$ | large |
| M0 | M1M2 | $-25.51$ | $< 10^{-300}$ | $-0.747$ | large |
| M0 | M6M7h | $+18.84$ | $< 10^{-300}$ | $+0.735$ | large |
| M0 | M4M5 | $-18.75$ | $< 10^{-300}$ | $-0.550$ | large |
| M3 | M4M5 | $+17.22$ | $< 10^{-300}$ | $+0.753$ | large |
| M4M5 | M6M7e | $-13.98$ | $< 10^{-300}$ | $-0.777$ | large |
| M1M2 | M4M5 | $+10.31$ | $< 10^{-300}$ | $+0.451$ | medium |
| M1M2 | M6M7e | $-4.04$ | $1.6 \times 10^{-4}$ | $-0.094$ | negligible |
| M1M2 | M3 | $-3.87$ | $2.2 \times 10^{-4}$ | $-0.045$ | negligible |
| M3 | M6M7e | $-0.98$ | $0.327$ | $-0.177$ | small |

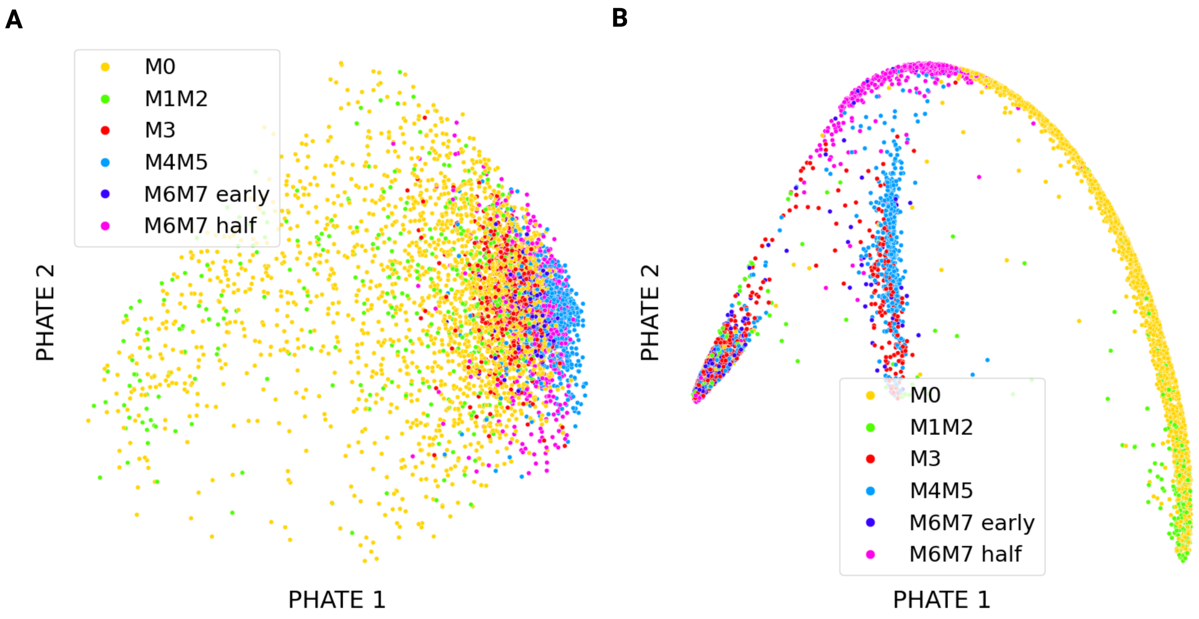


**Fig. S2.** PHATE embeddings of baseline feature matrices colored by cell-cycle stage, for comparison with Fig. 4C. (**A**) SHE features yield a relatively unstructured distribution in which cell-cycle stages are not clearly resolved. (**B**) CP features form a pronounced arc in which stages are ordered along the trajectory, recovering a cyclic organization similar to that seen with MT, though less tightly structured.

*Research*, 18(17):1–5, 2017.
26. Kaiming He, Xiangyu Zhang, Shaoqing Ren, and Jian Sun. Deep residual learning for image recognition. In *Proceedings of the IEEE Conference on Computer Vision and Pattern Recognition (CVPR)*, pages 770–778, 2016.
27. Matthias Feurer, Aaron Klein, Katharina Eggensperger, Jost Springenberg, Manuel Blum, and Frank Hutter. Efficient and robust automated machine learning. In *Advances in Neural Information Processing Systems*, volume 28, pages 2962–2970, 2015.
28. Henry B. Mann and Donald R. Whitney. On a test of whether one of two random variables is stochastically larger than the other. *The Annals of Mathematical Statistics*, 18(1):50–60, 1947.
29. Bernard L. Welch. The generalization of 'student's' problem when several different population variances are involved. *Biometrika*, 34(1–2):28–35, 1947.
30. Frank J. Massey. The kolmogorov–smirnov test for goodness of fit. *Journal of the American Statistical Association*, 46(253):68–78, 1951.
31. William H. Kruskal and W. Allen Wallis. Use of ranks in one-criterion variance analysis. *Journal of the American Statistical Association*, 47(260):583–621, 1952.
32. Olive Jean Dunn. Multiple comparisons using rank sums. *Technometrics*, 6(3):241–252, 1964.
33. Sture Holm. A simple sequentially rejective multiple test procedure. *Scandinavian Journal of Statistics*, 6(2):65–70, 1979.

**Table S4.** Per-image feature extraction time (ms / image, single-threaded) for MT (tensors + derived quantities including irreducible tensors not used in this analysis), SHE, and CellProfiler, measured on a representative sample of 20 images per dataset (Apple M1 Pro, 32 GB RAM; $n = 20$, seed = 42). MT and SHE values are mean ± std across individually timed images. CellProfiler value is total batch wall-clock divided by 20 (single subprocess invocation; per-image std not available). SHE timings are at $l_{\max} = 16$ for all datasets. Deep-learning ResNet baselines require GPU hardware; training and inference times are not included as they depend on hardware configuration and are not directly comparable to CPU-based feature extraction.

| **Dataset** | **MT (ms/img)** | **SHE $l_{\max}$=16 (ms/img)** | **CellProfiler (ms/img)** |
|---|---|---|---|
| Adrenal3D ($28^3$) | $\mathbf{28.6 \pm 14.2}$ | $33.9 \pm 10.1$ | 691 |
| Adrenal3D ($64^3$) | $148.4 \pm 43.0$ | $\mathbf{59.5 \pm 16.5}$ | 788 |
| Vessel3D ($28^3$) | $\mathbf{35.1 \pm 32.7}$ | $53.9 \pm 64.2$ | 1479 |
| Vessel3D ($64^3$) | $159.5 \pm 41.6$ | $\mathbf{60.6 \pm 12.2}$ | 1069 |
| hiPSC Nuclei ($\sim 200^3$) | $649.4 \pm 253.7$ | $\mathbf{250.7 \pm 41.5}$ | 2514 |

**Table S5.** Best LightGBM classification performance (balanced accuracy, BA; geometric mean, GM) per method and dataset at $28^3$ voxel resolution, mirroring Table 2 for the SVM classifier. Each cell shows mean ± std across three independent runs (each with a full independent optimize+evaluate cycle) and the best-performing feature subset. Row labels and abbreviations as in Table 2. **Bold** marks the highest value in each column. Deep-learning baselines (ResNet, AutoSklearn) are not repeated here; see Table 2.

| | **Adrenal3D** ($28^3$) | | **Vessel3D** ($28^3$) | | **Vessel3D** rand. orientation ($28^3$) | | **hiPSC Nuclei** | |
|---|---|---|---|---|---|---|---|---|
| **Method** | BA↑ | GM↑ | BA↑ | GM↑ | BA↑ | GM↑ | BA↑ | GM↑ |
| MT | **0.818 ± 0.006**<br>EVals+$\beta$+Scalars+Euler | **0.818 ± 0.006**<br>EVals+$\beta$+Scalars+Euler | **0.846 ± 0.012**<br>Tens.+Tr. | **0.846 ± 0.012**<br>Tens.+Tr. | **0.817 ± 0.013**<br>Invariants | **0.814 ± 0.015**<br>Invariants | **0.869 ± 0.004**<br>Tens.+Tr.+EVals | **0.868 ± 0.005**<br>Tens.+Tr.+EVals |
| SHE | 0.663 ± 0.033<br>Raw ($l$=14) | 0.699 ± 0.019<br>Raw ($l$=5) | 0.678 ± 0.032<br>Raw ($l$=6) | 0.664 ± 0.011<br>Raw ($l$=4) | 0.628 ± 0.023<br>Raw ($l$=1) | 0.592 ± 0.022<br>Raw ($l$=1) | 0.813 ± 0.005<br>Raw ($l$=11) | 0.799 ± 0.005<br>Raw ($l$=11) |
| CP | 0.777 ± 0.024<br>All features | 0.773 ± 0.005<br>All features | 0.789 ± 0.004<br>All features | 0.802 ± 0.016<br>All features | 0.765 ± 0.005<br>All features | 0.760 ± 0.006<br>All features | 0.778 ± 0.005<br>All features | 0.774 ± 0.008<br>Shape only |


34. Norman Cliff. Dominance statistics: Ordinal analyses to answer ordinal questions. *Psychological Bulletin*, 114(3):494–509, 1993.
35. Jacob Cohen. *Statistical Power Analysis for the Behavioral Sciences.* Lawrence Erlbaum Associates, Hillsdale, NJ, 2nd edition, 1988.
36. Pauli Virtanen, Ralf Gommers, Travis E. Oliphant, Matt Haberland, Tyler Reddy, David Cournapeau, Evgeni Burovski, Pearu Peterson, Warren Weckesser, Jonathan Bright, Stéfan J. van der Walt, Matthew Brett, Joshua Wilson, K. Jarrod Millman, Nikolay Mayorov, Andrew R. J. Nelson, Eric Jones, Robert Kern, Eric Larson, C. J. Carey, İlhan Polat, Yu Feng, Eric W. Moore, Jake VanderPlas, Denis Laxalde, Josef Perktold, Robert Cimrman, Ian Henriksen, E. A. Quintero, Charles R. Harris, Anne M. Archibald, Antônio H. Ribeiro, Fabian Pedregosa, Paul van Mulbregt, and SciPy 1.0 Contributors. SciPy 1.0: Fundamental algorithms for scientific computing in Python. *Nature Methods*, 17(3):261–272, 2020. doi: 10.1038/s41592-019-0686-2.

**Table S6.** 3D ResNet classification performance (balanced accuracy, BA; geometric mean, GM; accuracy, Acc; AUC) on the Allen Cell hiPSC nuclei dataset across three independent training runs. Values are mean $\pm$ std. **Bold** marks the highest value in each column. Results from 3D ResNet models trained in this work (see Supplementary Methods), adapted from the AllenCell `image_classifier_3d` framework (2).

| **Model** | BA↑ | GM↑ | Acc↑ | AUC↑ |
|---|---|---|---|---|
| ResNet-34 | **0.818 ± 0.007** | **0.782 ± 0.025** | **0.886 ± 0.003** | **0.980 ± 0.002** |
| ResNet-50 | 0.799 ± 0.010 | 0.767 ± 0.025 | 0.872 ± 0.008 | 0.977 ± 0.004 |
| ResNet-101 | 0.794 ± 0.022 | 0.762 ± 0.042 | 0.870 ± 0.014 | 0.976 ± 0.003 |

**Table S7.** Best SVM classification performance (balanced accuracy, BA; geometric mean, GM) per method and dataset at $64^3$ voxel resolution, mirroring Table 2 ($28^3$). Each cell shows mean $\pm$ std across three independent runs and the best-performing feature subset. Row labels and abbreviations as in Table 2. **Bold** marks the highest value in each column. Deep-learning ResNet baselines are evaluated at $28^3$ only (Table 2).

| | **Adrenal3D** ($64^3$) | | **Vessel3D** ($64^3$) | | **Vessel3D** rand. orientation ($64^3$) | |
|---|---|---|---|---|---|---|
| **Method** | BA↑ | GM↑ | BA↑ | GM↑ | BA↑ | GM↑ |
| MT | **0.852 ± 0.010**<br>Tens.+Tr.+EVals | **0.857 ± 0.005**<br>Tens.+Tr.+EVals | 0.824 ± 0.011<br>Tens.+Tr. | 0.831 ± 0.011<br>Tens.+Tr. | **0.786 ± 0.009**<br>Invariants | **0.788 ± 0.009**<br>EVals+$\beta$ |
| SHE | 0.728 ± 0.010<br>Raw ($l$=6) | 0.725 ± 0.010<br>Raw ($l$=6) | 0.713 ± 0.002<br>Raw ($l$=6) | 0.707 ± 0.006<br>Raw ($l$=6) | 0.640 ± 0.007<br>Raw ($l$=6) | 0.639 ± 0.008<br>Raw ($l$=6) |
| CP | 0.789 ± 0.007<br>All features | 0.788 ± 0.006<br>All features | **0.833 ± 0.016**<br>All features | **0.833 ± 0.016**<br>All features | 0.730 ± 0.021<br>SA+Solidity | 0.731 ± 0.020<br>SA+Solidity |

**Table S8.** Best LightGBM classification performance (balanced accuracy, BA; geometric mean, GM) per method and dataset at $64^3$ voxel resolution, mirroring Table S5 ($28^3$). Each cell shows mean $\pm$ std across three independent runs and the best-performing feature subset. Row labels and abbreviations as in Table 2. **Bold** marks the highest value in each column.

| | **Adrenal3D** ($64^3$) | | **Vessel3D** ($64^3$) | | **Vessel3D** rand. orientation ($64^3$) | |
|---|---|---|---|---|---|---|
| **Method** | BA↑ | GM↑ | BA↑ | GM↑ | BA↑ | GM↑ |
| MT | **0.789 ± 0.011**<br>Invariants | **0.788 ± 0.011**<br>Invariants | **0.833 ± 0.014**<br>Tens.+EVals | **0.832 ± 0.014**<br>Tens.+EVals | **0.832 ± 0.009**<br>Invariants | **0.831 ± 0.009**<br>Invariants |
| SHE | 0.697 ± 0.019<br>Raw ($l$=7) | 0.697 ± 0.018<br>Raw ($l$=7) | 0.670 ± 0.023<br>Raw ($l$=3) | 0.663 ± 0.029<br>Raw ($l$=6) | 0.601 ± 0.007<br>Raw ($l$=1) | 0.577 ± 0.006<br>Raw ($l$=1) |
| CP | 0.786 ± 0.006<br>All features | 0.787 ± 0.008<br>All features | 0.794 ± 0.006<br>All features | 0.788 ± 0.013<br>All features | 0.762 ± 0.015<br>All features | 0.775 ± 0.014<br>All features |
